# Adaptation of *Burkholderia cenocepacia* to low oxygen drives changes consistent with adaptation to chronic infection

**DOI:** 10.1101/2025.08.08.669301

**Authors:** Ciarán J. Carey, Joanna Drabinska, Niamh Duggan, Siobhán McClean

## Abstract

**Background:** Cystic fibrosis (CF) is characterised by chronic respiratory infections, involving opportunistic pathogens, including *Burkholderia cenocepacia*. The CF lung comprises hypoxic niches that drives bacterial adaptation and the adaptability of pathogens to this environment is key to their successful colonisation. We previously identified several proteins encoded on a low-oxygen activated (Lxa) locus that were significantly increased in abundance in late chronic infection *B. cenocepacia* isolates. However, the impact of long-term hypoxia exposure on *B. cenocepacia* adaptation remains unclear.

**Results:** To investigate the role of hypoxia in driving traits associated with chronic infection, we exposed an early infection *B. cenocepacia* isolate to low (6% O₂) or atmospheric oxygen (21% O₂) over 22 days. By day 22, 364 proteins were significantly increased in abundance in hypoxia-adapted cultures relative to the ancestral strain. Overall, 1066 individual proteins were significantly increased in abundance in the hypoxia-adapted cultures relative to normoxia-adapted cultures, across four different timepoints from day 1 to day 22. Comparative proteome analysis identified 81 proteins with consistent changes in abundance both in hypoxia-adapted cultures and the respective late infection isolate relative to the ancestral strain (the early infection isolate), including lxa-encoded proteins and the FixK transcriptional regulator. Proteins associated with shikimate pathways were also significantly changed in abundance. Importantly, hypoxia-adapted cultures showed increased survival in CF macrophages, increased attachment to CF lung cells, elevated protease activity, greater resistance to ceftazidime and ciprofloxacin, all of which are consistent with adaptations observed in late chronic infection isolates. Hypoxia-adapted cultures also displayed enhanced virulence in *Galleria mellonella* larvae, as did the late infection isolate.

**Conclusions:** The changes in phenotype and proteome of *B. cenocepacia* observed after long-term hypoxia suggest that hypoxia may drive the adaptation to chronic infection, promoting survival in macrophages, host-cell attachment, antibiotic resistance and protease activity. Therapeutic strategies that modulate oxygen availability or target hypoxia-sensing may hold promise in preventing or mitigating chronic infection in CF.

## Introduction

Cystic fibrosis (CF) is a genetic disorder caused by mutations in the Cystic Fibrosis Transmembrane Conductance Regulator (CFTR) protein [1]. This results in impaired chloride ion transport that ultimately creating conditions that promote chronic bacterial and fungal infections that are difficult to treat — a hallmark of cystic fibrosis [2, 3]. The inherent adaptability and plasticity of opportunistic pathogens, such as *Pseudomonas aeruginosa* or *Burkholderia cepacia* complex (Bcc) facilitate the transition from acute to chronic infection [4]. Exposure to antibiotics and other environmental stressors, force pathogens to adapt in order to survive and thrive [5]. The response to these exogenous stimuli are often shared, and ultimately dictate bacterial behaviours [6]. Therefore, adaptation to different environmental conditions can have significant and often unforeseen consequences for the host, particularly in the context of chronic infection [7, 8].

Bcc consists of 26 species [9], of which *B. cenocepacia* is the most virulent for people with CF [10, 11]. *B. cenocepacia* is a rod-shaped, Gram-negative, opportunistic pathogen often found in soil and water environments [12]. It is inherently antibiotic resistant and harbours a multireplicon genome with an exquisite capacity for pathogenicity [13]. Chronic infections with Bcc are often associated with rapid declines in patient lung function, significantly increasing morbidity and mortality in individuals with CF [14, 15].

The CF lung environment is host to a variety of environmental pressures, requiring opportunistic pathogens such as *B. cenocepacia* to adapt effectively in order to successfully colonise [16]. Mucus plugging and immune cell function, as a result of a sustained immune response, contribute to the development of hypoxic niches in the lung, and pulmonary exacerbations arising from persistent infection further reduce oxygen availability and create severe oxygen gradients [17]. Successful adaptation to these low oxygen niches contributes to the development of chronic infection [18]. Sass et al., (2013) discovered a 50 gene cluster dubbed the ‘low-oxygen activated’ (lxa) locus after comparative transcriptional analysis of a *B. cenocepacia* strain exposed to low oxygen for two hours [19]. We subsequently identified significant increases in the abundance of 149 and 189 proteins respectively, in the proteomes of two late infection sequential CF isolates relative to their respective early infection counterparts respectively, and the same subgroup of 19 Lxa-encoded proteins showed increased abundance in both late infection isolates [20]. This indicates that both late infection isolates adapted over time, and the consistent effect on Lxa-encoded proteins highlights a potential role of hypoxia in the adaptive process.

Following that, the aim of this study was to investigate whether low oxygen availability in the CF lung drives mechanisms of adaptation of *B. cenocepacia*. We exposed one of the early infection *B. cenocepacia* isolates (P1E) to either low oxygen (6% oxygen) or normoxia (21% oxygen) for up to 22 days and examined the changes in proteome and phenotype in response to hypoxia over the extended period. We focussed on the proteome because this is more directly linked to alterations in phenotype and allowed us to compare with our previous work. Moreover, stable changes in proteome are less well investigated than those of the genome in adaptation studies. Consequently, this will provide new insights into the mechanisms involved in adaptation to low oxygen and the establishment of chronic infection.

## Materials and Methods

### Bacterial Strains

*B. cenocepacia* early (P1E) and late (P1L) sequential clinical isolates were previously characterised (Cullen et al., 2018). Overnight *B. cenocepacia* cultures were routinely grown in lysogeny broth (LB) (Sigma-Aldrich, St. Louis, MO, USA) with orbital agitation (200 rpm) at 37°C, 5% CO_2_, unless stated otherwise.

### Long term Exposure of Bcc Strains to Hypoxia

Three 100 ml flasks containing 30ml of LB broth, were pre-equilibrated for 18 hours in 6% O_2_ and 5% CO_2_ at 37°C in O_2_ Control InVitro Glove Box (PHGB model, Coy Laboratory Products, MI, United States) and another three flasks containing 30ml of LB broth, were pre-equilibrated for 18 hours at 21% O_2_ and 5% CO_2_ in the Thermo Scientific SteriCycle CO_2_ Incubator Model 371 (Thermo Fisher Scientific, Waltham, MA, USA). Each flask was inoculated with 2 ml of an overnight *B. cenocepacia* P1E culture at a density of OD_600nm_ 0.1 and cultured for up to 22 days in hypoxia (6% O_2_) or normoxia (21% O_2_) respectively at 200rpm in 37°C, 5% CO_2_. The cultures were sub-cultured into fresh LB every 48 hours by transferring three ml of each culture to pre-equilibrated flasks without removal from their respective environments and samples simultaneously harvested for proteome analysis, growth analysis (OD_600nm_ measurements and CFU counts) and glycerol stocks. The glycerol stocks (50%, LB (v/v)) were prepared and stored immediately for future phenotypic analyses, while, bacterial cells were pelleted at 12,000 *xg* for 2 min and stored at -80°C for proteome analyses. This process was repeated on every 48 h for 22 days. All sampling and sub-culturing was completed in hypoxic conditions (6% O_2_) or normoxic conditions (21% O_2_), as appropriate.

### Proteome Analysis

Proteome analysis was performed as previously described [20] with minor modifications. The samples were retrieved from storage at -80°C and whole cell lysates were resuspended in 1 ml of ice-cold lysis buffer containing 40mM Tris-HCl (pH 7.8) and 1x cOmplete™, EDTA-free protease inhibitor cocktail (Roche, Basel, Switzerland). The cells were placed on ice and disrupted by sonication with a sonic dismembrator and microtip 6mm probe (model 505; Fisherbrand, Thermo Fisher Scientific) for eight 30s bursts followed by 30s rest. Cell debris and any remaining intact cells were pelleted using centrifugation (10,000 *xg* for 30 min). Following sample preparation and dialysis, aliquots from each sample were placed in fresh tubes and samples were dried in a vacufuge concentrator system (model 5301; Eppendorf, Hamburg, Germany) at 45°C and resuspended in 20 μl of resuspension buffer containing 0.5% Trifluoroacetic acid (TFA) in MilliQ water. ZipTips (Merck Millipore, Burlington, MA, USA) were used to purify samples prior to analysis on a Bruker TimsTOF Pro mass spectrometer (Bruker, MA, USA) with Evosep One chromatography system (Evosep, Denmark). Peptides (1-2 μg) were separated on a 8 cm analytical C18 column at the pre-set rate of 33 samples per day. A trapped ion mobility (TIMS) analyser was synchronized with a quadrupole mass filter with acquisition rates of 100 Hz; accumulation and ramp times of 100 ms.; and ion mobility (1/k0) range from 0.62 to 1.46 Vs/cm. Spectra were recorded in the mass range from 100 to 1,700 m/z.

### Determination of Antibiotic Susceptibility

Antibiotic susceptibility was measured using the Kirby-Bauer method on Mueller Hinton agar (Neogen, Lansing, MI, USA) plates. Mid-log phase cultures adjusted to McFarland standard 1 and plated on 145mm plates. Each plate was inoculated with 350 μl of each adjusted culture and spread using a cotton swab, rotating the plates 5 times to ensure homogeneity. Antibiotic discs (Oxoid, Hampshire, UK) were then added using a sterile forceps and plates incubated for 48 hours at 37°C and 5% CO_2_. Zones of inhibition were measured using a ruler after 2 days of growth, an mean was used for statistical analysis.

### Determination of Bacterial Protease Activity

Protease activity was measured on 1% milk agar plates (145 mm). A 1:10 (w/v) of powdered milk and dH_2_O was sterilised at 110°C for 10 minutes before being added to sterile LB agar. Plates were inoculated with mid-log phase cultures (1 μl) which had been adjusted to McFarland standard of 1 and incubated for 48 hours at 37°C and 5% CO_2_. Zones of clearance were measured by two perpendicular measurements after 2 days of growth, an average was used for statistical analysis.

### Analysis of protease profile by zymography

A 50 ml overnight *B. cenocepacia* culture was harvested and centrifuged at 5,000 *xg* for 10 minutes at 4°C. The bacterial supernatant was then concentrated using Amicon Ultra-15 centrifugal filters (Sigma-Aldrich, St. Louis, MO, 32USA) by centrifugation in an angled rotor centrifuge at 6,000 *xg*. The concentrated protein was stored at 4°C until required (or at -80°C for long term storage) and the pellet discarded. The protein concentration was measured with Bradford reagent (Thermo Fisher Scientific) and the concentrations adjusted accordingly. Bovine gelatin (1 mg/ml, Sigma-Aldrich) was dissolved in dH_2_O at 37°C before addition to a resolving gel solution for the zymogram. A standard SDS-PAGE gel was prepared in parallel. The protein samples were heat-denatured and prepared for loading on an SDS-PAGE gel by mixing the protein samples with 5x Laemmli Buffer (10% (v/v) glycerol, 2% (w/v) SDS, 62.5mM Tris-HCl (pH 6.8), 0.01% (w/v) bromophenol blue) at a ratio of 5:1. Precision Plus Protein Dual Colour Standard protein ladder (12 μl) (BioRad, Hercules, CA, USA) was loaded and PA01 supernatants were used as positive control. The gels were run at 110V for 100 minutes before rinsing in dH_2_O. The SDS was removed by incubation in 2.5% Triton X-100 in PBS, with two changes of buffer following each 30-minute incubation period, with agitation. This was followed by three dH_2_O washes. The zymogram was then incubated in digestion buffer (25mM Tris base, 192mM glycine, 50 mM Tris-HCl, 1 M Tris-HCL, 1 mM CaCl2,10 mM ZnCl2,1 mM DTT, 500 ml with dH_2_O) overnight at 37°C. Both gels were stained with Coomassie blue (VWR) before destaining and analysis.

### Biofilm Formation

Biofilm formation was determined using crystal violet staining, as previously described [21]. Round-bottom 96-well plates were inoculated with overnight *B. cenocepacia* cultures adjusted to OD_600nm_ 0.1 in fresh, pre-warmed LB broth and 100 µl of each diluted culture was transferred into 8 wells. The 96-well plates were incubated statically at 37°C and 5% CO_2_ for 24, 48 or 72 h and stained with crystal violet 0.1% w/v before the OD_600nm_ was measured with a plate reader.

### Determination of Host-cell Attachment

Attachment to CF lung epithelial cells was performed as previously described [20] with minor modifications. CFBE41o^-^ cells were seeded in 24-well plates coated with minimal essential media (MEM) containing 10% w/v bovine serum albumin (BSA), 1% v/v collagen and 1% v/v fibronectin at a density of 4 x 10^5^ cells/well in antibiotic-free MEM and incubated at 37°C and 5% CO_2_ overnight. *B. cenocepacia* cultures were grown to mid-logarithmic phase in LB broth, resuspended in MEM and added to each corresponding well at a concentration of 4 x 10^6^ CFU/well (MOI 10:1). The 24-well plates were then centrifuged at 252 *xg* for five minutes followed by a 30-minute incubation at 37°C, 5% CO_2_ to allow for bacterial attachment. The wells were then washed with PBS to remove any unadhered cells, before 500 µl of lysis buffer (0.25 % Triton X-100 in PBS) was added for 20 minutes at room temperature. Following enumeration, the CFU/ml was determined, and the percent attachment was calculated as described previously [20].

### CF PBMC-derived Intramacrophage Survival

CF PBMCs were derived as previously described [21] and seeded at a concentration of 1 x 10^5^ cells/ml in 24-well plates and incubated overnight. *B. cenocepacia* cultures were grown to mid-logarithmic phase in LB broth, resuspended in RPMI-1640 medium and added to each corresponding well at a concentration of 5 x 10^5^ CFU/well (MOI 5:1). The plates were centrifuged at 1100 *xg* for five minutes and then incubated at 37°C, 5% CO_2_ for two hours to allow for uptake. The cells were incubated with antibiotics for two hours at 37°C, 5% CO_2_ to kill any extracellular bacteria. Cells were lysed with 0.25% v/v Triton X-100 for 20 minutes at RT, at time zero and at 24 h. Following enumeration, the CFU/ml were determined, and the percentage attachment was calculated as described previously [21].

### Tryptophan Supplementation Assay

96-well plates were inoculated with overnight *B. cenocepacia* cultures and washed with PBS twice before being adjusted to OD_600nm_ 0.1 in M9 minimal media, and 150 µl of each diluted culture were transferred into eight wells. Six wells were supplemented with 100mM tryptophan (Thermo Scientific) and the 96-well plates incubated at 37°C with shaking for 24 hr. Optical density (OD_600nm_) was measured with a Syngery H1 microplate reader every 30 minutes.

### *Galleria mellonella* Acute Infection Model

*G. mellonella* wax moth larvae (deKammieshop, Waalwijk, NL) were maintained at 15°C for a minimum seven days post-delivery prior to use and used within four weeks. *B. cenocepacia* cultures were cultured to mid-logarithmic phase and resuspended in 1 ml PBS before being diluted to a range of concentrations (2 x 10^5^ to 2 x 10^9^ CFU/ml). Ten healthy, cream larvae weighing between 0.2 and 0.4g were selected and each strain was injected into the hindmost left proleg at a volume of 20 µl per dilution, using a sterile Terumo 0.3ml syringe (Terumo, Leuven, Belgium). The same volume of each dilution was plated on LB agar plates in duplicate for CFU enumeration. The larvae were then incubated at 37°C and the percentage survival of the larvae was measured at 24, 48 and 72 hours and graphed using Kaplan-Meier curves. The percentage larval survival was plotted against the CFU bioburden in order to calculate the LD30 value.

### Statistical analysis

All statistical analyses were performed using GraphPad Prism (v8) unless otherwise stated. All analyses were completed using one-way ANOVA following normalisation of the data, with the exception of the measurement of antibiotic resistance where a paired student t-test was used. Proteome data were analysed by one-way ANOVA for three-way comparisons and by t-tests for pairwise comparisons of samples (p < 0.05; fold change >1.5).

## Results

### Adaptation to Low Oxygen

In order to investigate the impact of long-term exposure to low oxygen on *B. cenocepacia*, we exposed the early infection isolate, P1E [22] to either low oxygen (6% O_2_) or atmospheric oxygen (21% O_2_) over a 22-day period (**Figure 1**). Three independent flasks were exposed to each condition and sub-cultured into fresh lysogeny broth (LB) and sampled every two days. Four different colony variants emerged in plated samples (**Figure 2**) over time across both sets of conditions. Over the course of 22 days, the typical *B. cenocepacia* colony variant (**Figure 2A**) remained the predominant variant. Small colony variants (SCVs) (**Figure 2B**) emerged in both conditions (**Figure 2C and 2D**). A ‘wrinkly spreader’ morphotype (**Figure 2E**) also emerged sporadically in both conditions but less frequently in normoxia. Interestingly, a novel morphotype with a ‘doughnut’ like appearance (**Figure 2F**) was observed exclusively in hypoxia-exposed cultures (**Figure 2G**), which was stable if subsequently cultured under normoxic conditions following multiple passages.

**Figure 1.**
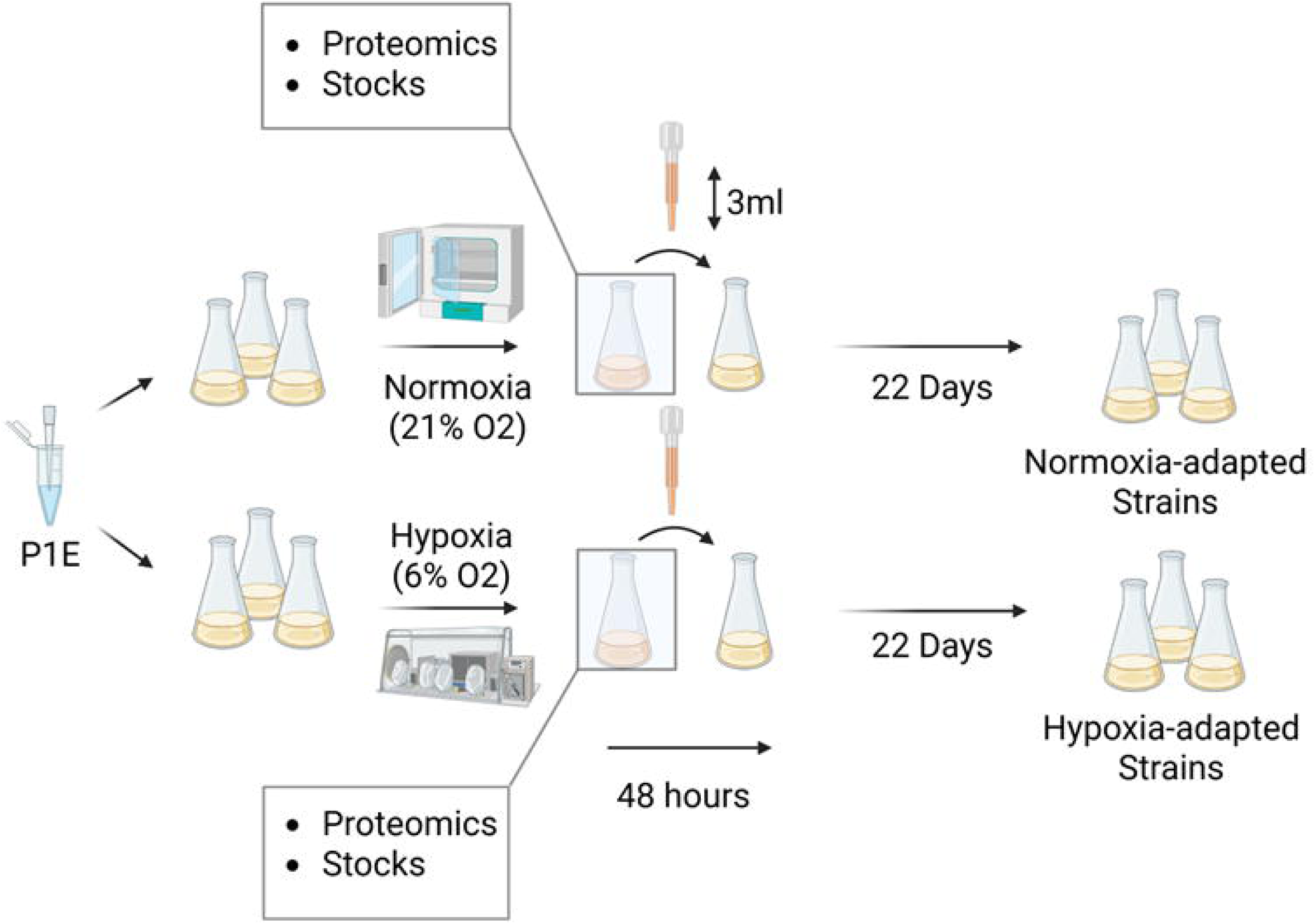
Experimental low oxygen adaptation study design over 22 days. Overnight cultures of early infection isolate (P1E) were inoculated int three flasks with either normoxia-or hypoxia-equilibrated medium and incubated with shaking in 21% or 6% oxygen. Every two days, aliquots (3 ml) were transferred to a fresh flask of equilibrated medium under normoxia or hypoxia as appropriate to avoid oxygen fluctuations. Samples were also removed for phenotyping and proteome analysis and stored for future use. Long-term exposure of an early *B. cenocepacia* clinical isolate to low oxygen or normal oxygen conditions was maintained over a 22-day period.

**Figure 2.**
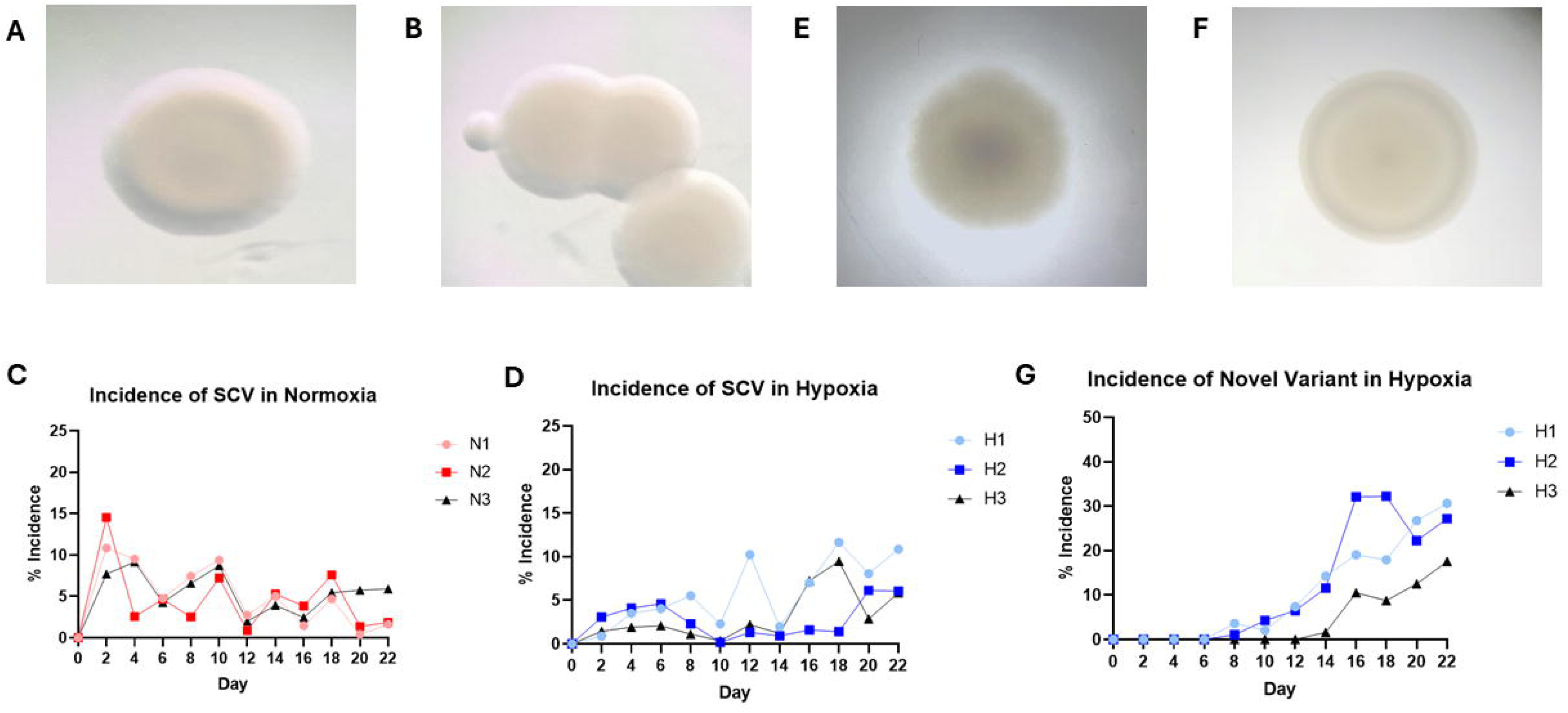
Impact of long-term exposure to hypoxia on colony morphology. Representative images of the different *B. cenocepacia* colony variants observed during the long-term adaptation study and their incidences. **A)** Normal colony variant. **B)** small colony variant (SCV) and its incidence in **C)** normoxia or **D)** hypoxia; **E)** wrinkly spreader variant; **F)** novel morphotype with a ‘doughnut’ like appearance and **G)** its incidence in hypoxia.

### Comparison of proteomes of long-term hypoxia- and normoxia-adapted cultures

In order to elucidate the changes in both the proteome of the hypoxia-adapted cultures (HACs) and the normoxia-adapted cultures (NACs) relative to the ancestral strain, P1E (Day 0), a longitudinal proteome analysis was conducted, by comparing the proteomes of the early infection isolate to the proteomes of the HACs and NACs at day 22. We analysed the proteomes of the adapted cultures, to focus on the impact of hypoxia on the expressed and post-translationally modified proteins that would directly impact phenotype.

A total of 364 proteins were significantly changed in abundance (p<0.05) in the proteomes of the HACs at day 22 relative to the ancestral strain on day 0, while 381 proteins were significantly changed in abundance in the proteomes of the NACs at day 22 relative to Day 0 (**Figure 3A**). Comparison of the proteomes of HACs at day 22 with those of NACs on the same day showed 232 proteins that were significantly changed in abundance. Culture conditions were identical between these two conditions apart from the oxygen availability, thus the change in the abundance of proteins is directly attributable to the difference in oxygen levels. To elucidate the potential biological impact of hypoxia exposure on the proteomes of the early infection isolate, eggNOG mapper and assigned GO terms were used (**Supplementary Table 1**). The functional enrichment analysis revealed 19 enriched categories including those associated with cellular processes and signalling, information processes and storage, and metabolism. The categories with the greatest number of proteins changed in abundance were those related to cell wall biogenesis, energy production and conversion and amino acid transport and metabolism.

**Figure 3.**
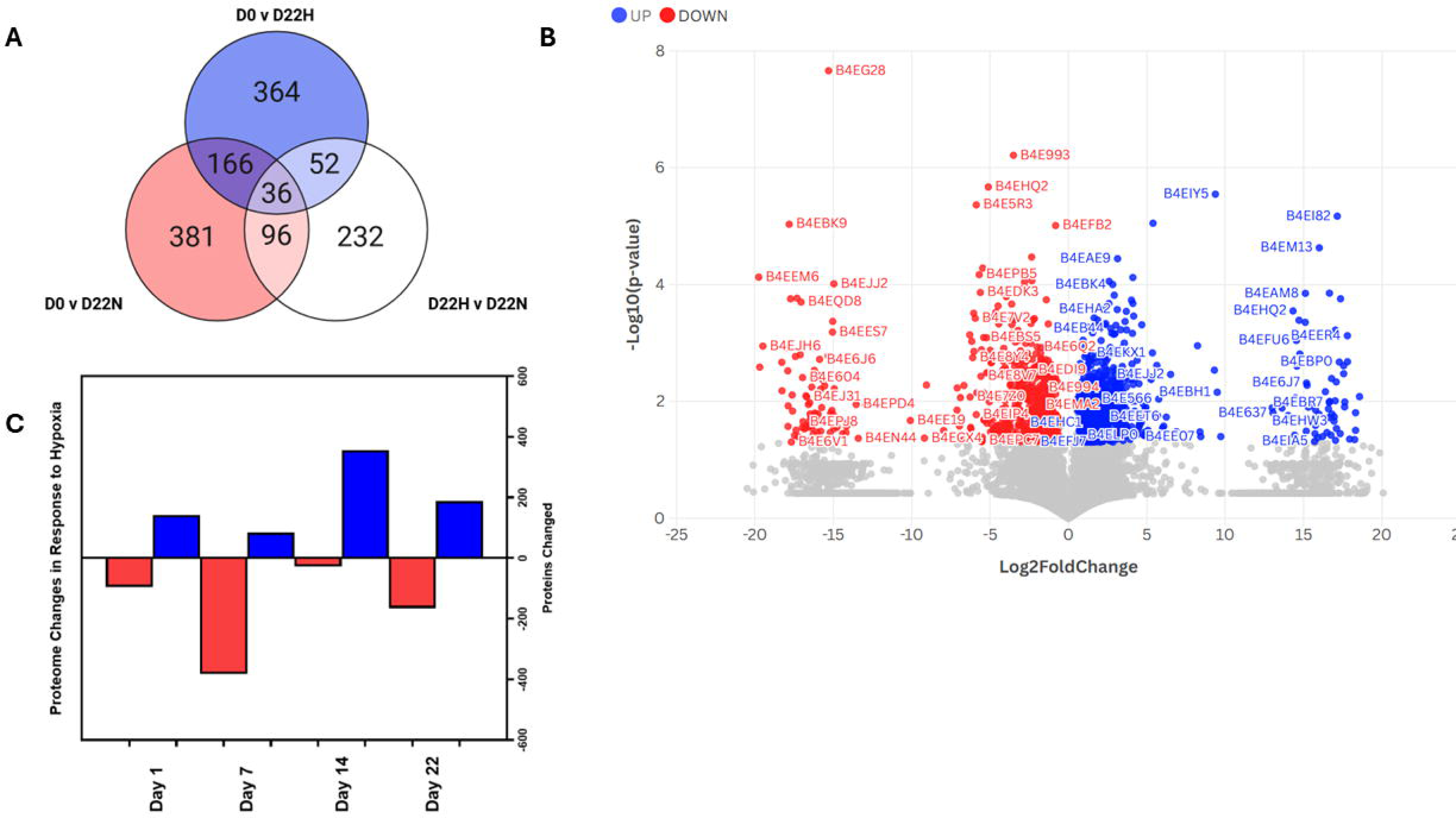
Changes in protein abundance in the proteome of the hypoxia-adapted ***B. cenocepacia* cultures relative to *B. cenocepacia* cultures. A)** Venn diagram illustrating the number of proteins significantly changed in abundance between the ancestral strain, P1E (D0) and HACs (D22H); ancestral strain (D0) and NACs (D22N) and between the HACs and the NACs in the longitudinal analysis. **B)** Volcano plot depicting log2 fold change in proteins significantly changed in abundance in the proteome of the hypoxia-adapted cultures relative to the normoxia-adapted cultures over the 22-day period. **C)** Differences in relative protein abundance in proteome hypoxia-adapted cultures relative to normoxia-adapted cultures; blue bars indicative of increased abundance, red bars indicative of decreased abundance.

To further investigate the impact of the adaptation to hypoxia over time, pairwise comparisons were conducted on the proteomes of the HACs relative to the NACs, at four different time-points to highlight proteins that were consistently increased or decreased in abundance over the time course. In total, 1066 individual proteins were significantly changed in abundance over the 22-day period (including 374 proteins that were changed in abundance on multiple days) by >1.5 fold or more (p<0.05) (**Figure 3B**). There were 237 proteins that were significantly changed in abundance in the HACs relative to the NACs on day 1, while by days 7, 14 and 22, there were 467, 383 and 353 proteins that were significantly changed in abundance in the HACs compared with the NACs (**Figure 3C**). Interestingly, 61 proteins were found to be unique to hypoxia exposed cultures and were reproducibly identified in all three independently adapted HAC flasks while being absent or undetectable in all three NAC flasks.

To investigate the impact of long-term exposure to hypoxia on bacterial adaptation, the day 22 proteome samples were analysed using the eggNOG mapper and assigned GO terms to determine their biological functions. The functional enrichment analysis revealed 20 enriched categories (**Supplementary Table 2**) including those associated with cellular processes and signalling, information processes and storage, and metabolism. There were significant changes in the abundance of proteins related to translation and transcription, replication, energy metabolism, amino acid and organic compound transport, in addition to secondary metabolite biosynthesis and processing. Interestingly, the abundance of proteins involved in host defence, intracellular trafficking, antibiotic resistance and cell wall biogenesis were also altered. There were 22 proteins identified that were unique to all three HACs on day 22 and not detected in NACs. In contrast, 16 proteins were uniquely identified in all NACs on day 22, highlighting the distinct differences in adaptation to each condition (**Supplemental Table 3**).

### Hypoxia Drives Changes Associated with Increased Tolerance to Oxidative Stress

At least ten proteins implicated in oxidative stress tolerance significantly increased in abundance in the proteome of the HACs relative to the early infection isolate (**Table 1**), including multiple universal stress proteins (USPs), heat shock protein (HSPs), cold shock proteins (CSPs), alkyl hydroperoxide reductases, thioredoxins and short chain dehydrogenases. Interestingly, SodB, a superoxide dismutase, was increased in abundance ∼2.5 fold in the proteome of the HACs relative the early infection isolate. The flavohemoprotein, WbrA, was also significantly increased in abundance by ∼15 fold in the HACs after 22 days. In the pairwise comparison of the HACs and the NACs, at least 15 proteins associated with enhanced tolerance to oxidative stress were revealed to be significantly increased in abundance in the HACs relative to the NACs. Another flavohemoprotein, HmpA, was significantly increased in abundance by ∼300 to 350-fold on days 14 and 22 (**Table 2**). A thiol-disulfide oxidoreductase, which is crucial in the detoxification of reactive oxygen species [23], was unique to HACs on day 22. Significantly, AhpC, which is associated with intracellular survival [24], was significantly increased in abundance in the HACs on days 14 and 22 by ∼2 to ∼3.5 fold while AhpD was increased on days 1 and 22 (∼2 to 4-fold). A putative thioredoxin was significantly increased in abundance in the HACs by ∼2 to ∼5.5 fold on days 7 and 14, while a thioredoxin reductase was increased on the same days (2.5 to 3 fold). Other redox proteins, e.g. a putative thiol peroxidase and a NADH-dependent flavin oxidoreductase were also significantly increased on days 14 and day 22. Furthermore, proteins associated with cysteine biosynthesis, cysteine modification and cysteine metabolism were significantly increased in abundance in the HACs relative to the NACs.

**Table 1.** Examples of proteins significantly changed in abundance in abundance in the proteome of the day 22 hypoxia-adapted cultures relative to early infection isolate.

| Protein ID | Gene ID (BCA) | Protein Function | Comparison | Fold Change | <sup>a</sup>UP | <sup>b</sup>MW | <sup>c</sup>SC |
| --- | --- | --- | --- | --- | --- | --- | --- |
| **Oxidative stress** |  |  |  |  |  |  |  |
| B4E9F6 | L2757 | Superoxide Dismutase SodB | D22H v D0 | 2.56 | 13 | 21.128 | 78.1 |
| B4EH97 | M1217 | Alkyl hydroperoxide reductase C | D22H v D0 | 1.81 | 7 | 20.728 | 46.5 |
| B4EC21 | L2014 | Alkyl hydroperoxide reductase D | D22H v D0 | 2.61 | 11 | 18.766 | 56.6 |
| B4ECT7 | L3146 | Chaperonin GroEL | D22H v D0 | 2.4 | 62 | 56.98 | 96 |
| B4ECT8 | I3147 | Co-chaperonin GroES | D22H v D0 | 1.68 | 16 | 10.49 | 95.9 |
| B4EMN5 | M0833 | OsmC-like protein | D22H v D0 | 8.01 | 5 | 14.673 | 27.1 |
| B4EJJ0 | M1500 | Universal Stress Protein | D22H v D0 | 12.05 | 25 | 33.296 | 68.5 |
| B4EG36 | M0050 | Universal Stress Protein A | D22H v D0 | 3.39 | 17 | 16.473 | 96.1 |
| B4EDZ2 | L3270 | Heat Shock Protein 70 DnaK | D22H v D0 | 1.46 | 69 | 69.741 | 83.2 |
| B4E730 | L2464 | Short chain dehydrogenase | D22H v D0 | 181.5 | 12 | 24.961 | 66.7 |
| B4EBR9 | L3035 | Thioredoxin reductase TrxB | D22H v D0 | 2.49 | 20 | 34.35 | 78.4 |
| **Aromatic Amino Acid Metabolism** |  |  |  |  |  |  |  |
| B4E7H2 | L1467 | Chorismate synthase AroC | D22H v D0 | 3.38 | 11 | 38.908 | 36.3 |
| B4E596 | L0210 | TetR family regulatory protein PaaR | D22H v D0 | 1.85 | 8 | 23.022 | 45.3 |
| B4EDA1 | L2179 | Enolase | D22H v D0 | 1.79 | 25 | 45.686 | 65.6 |
| B4E568 | L0011 | AsnC family regulatory protein | D22H v D0 | 4.55 | 2 | 17.78 | 21.1 |
| B4ELC9 | M1751 | Isochorismatase | D22H v D0 | 5.39 | 5 | 19.767 | 43.9 |
| **Pathogenesis** |  |  |  |  |  |  |  |
| B4E7G7 | L1046 | Phospholipase C PlcN | D22H v D0 | 9.45 | 2 | 21.1 | 13.1 |
| B4E5Y8 | L0264 | Delta-aminolevulinic acid dehydratase HemB | D22H v D0 | 2.79 | 17 | 37.102 | 62 |
| B4EI91 | M0280 | Phospholipid-binding protein | D22H v D0 | 6.31 | 11 | 23.517 | 60.2 |
| B4EF89 | M0944 | Lipoprotein | D22H v D0 | 2.71 | 7 | 25.194 | 43.4 |
| B4EPE7 | S0759 | Peptidoglycan-binding membrane protein | D22H v D0 | 6.22 | 4 | 484.63 | 1 |
| **Antibiotic Resistance** |  |  |  |  |  |  |  |
| B4EKD7 | M2652 | TetR family regulatory protein | D22H v D0 | 26420 | 2 | 20.448 | 16.5 |
| B4E6J6 | L3466 | UDP-N-acetylmuramoyl-tripeptide--D-alanyl-D-alanine ligase MurF | D22H v D0 | 2.52 | 17 | 48.154 | 52.6 |
| B4EBM6 | L1999 | MarR family regulatory protein | D22H v D0 | 1.95 | 5 | 16.676 | 36.2 |
| B4EPN0 | S0113 | ABC transporter protein | D22H v D0 | 5.58 | 1 | 29.075 | 4.9 |
| **DNA Repair** |  |  |  |  |  |  |  |
| B4ED83 | L2160 | DNA mismatch repair protein MutS | D22H v D0 | -2.92 | 18 | 95.986 | 25.5 |
| B4E6I0 | L2454 | Topoisomerase IV subunit A ParC | D22H v D0 | -1.78 | 21 | 84.107 | 30.1 |
| B4E6I1 | L2455 | Topoisomerase IV subunit B ParE | D22H v D0 | -3.03 | 19 | 72.669 | 33 |

B.c: *Burkholderia cenocepacia*, <sup>a</sup>UP: Unique peptides: The number of peptide sequences unique to a protein, <sup>b</sup>MW: Molecular weight as determined by TIMS-ToF LC-MS using Burkholderia cenocepacia J2315 database, <sup>c</sup>SC: % sequence covered by matching peptides for the identified protein in the database

**Table 2.**
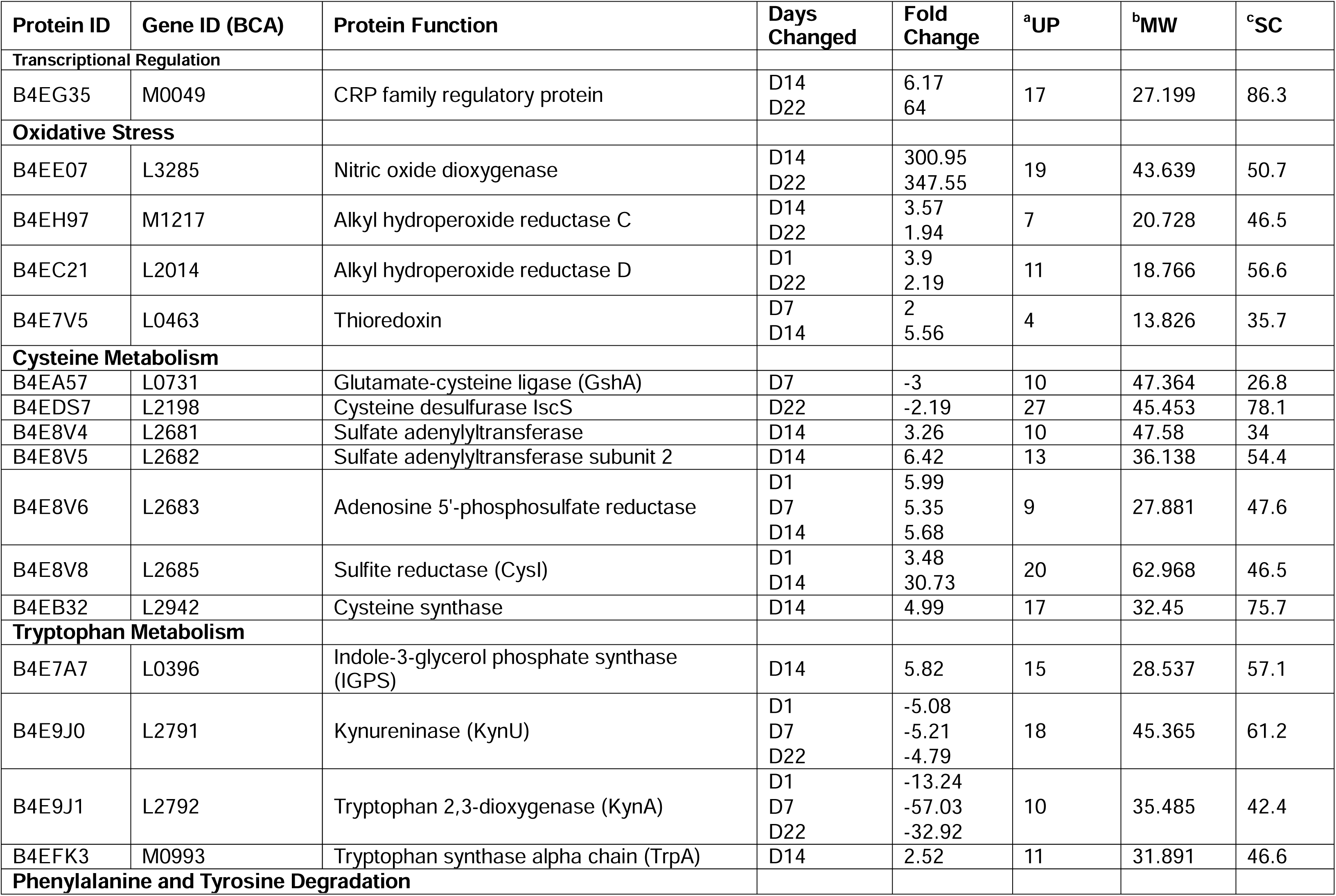
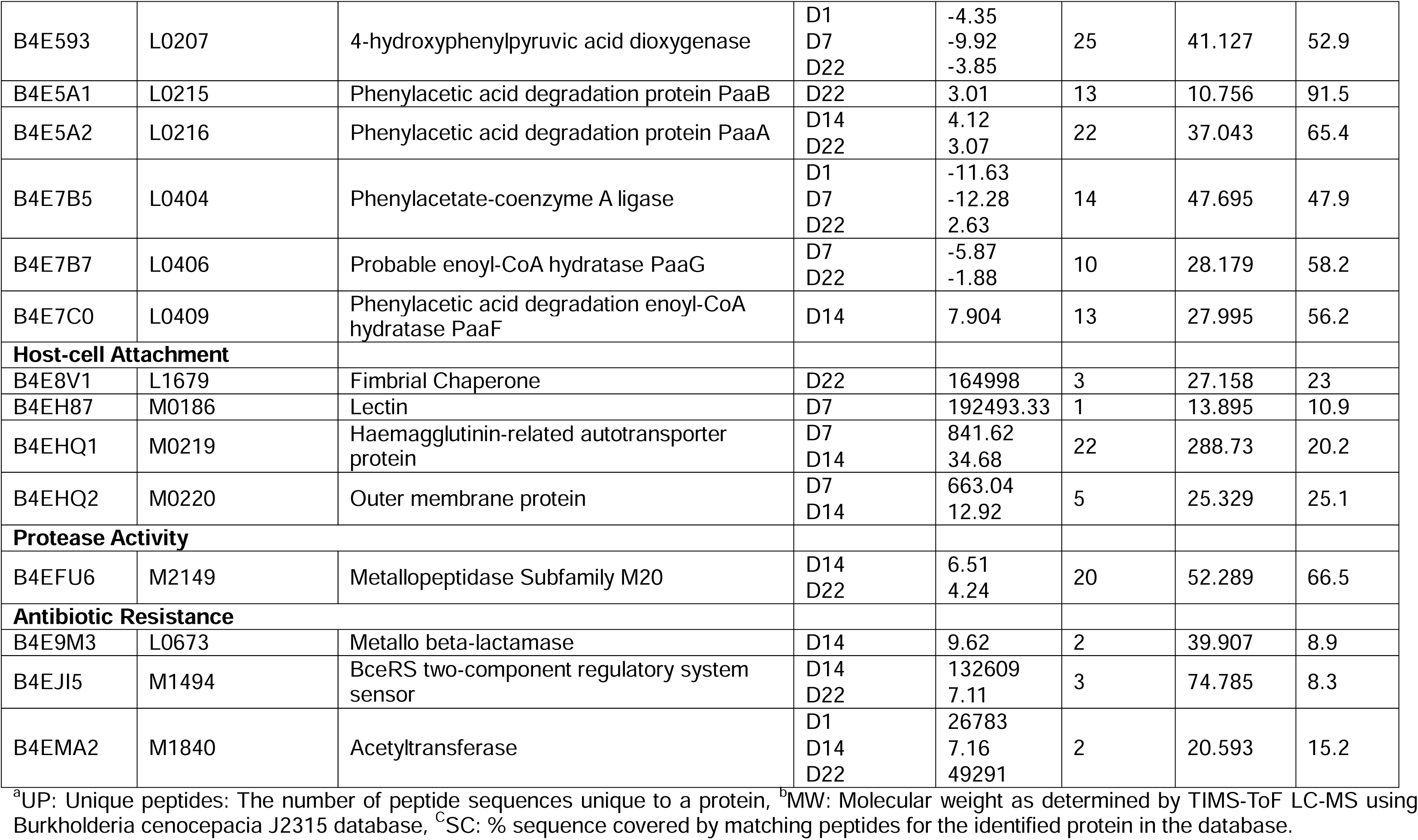
Examples of proteins significantly changed in abundance in hypoxia-adapted cultures relative to normoxia-adapted cultures.

Cysteine plays a crucial role in the response to reactive oxygen speces (ROS) [23, 25]. Proteins associated with cysteine biosynthesis were significantly increased in abundance in the HACs relative to NACs, with an O-acetylhomoserine thiol lyase uniquely identified in HACs on Day 7 (**Table 2**). CysI, a putative sulfite reductase was significantly increased in abundance in the HACs by ∼3.5 fold on day 1 and by ∼30.5 fold on day 14 while CysH, an adenosine 5’-phosphosulfate reductase also encoded by the *cys* regulon, was increased by 5.5 to 6 fold on days 1, 7 and 14. Additional proteins encoded by the *cys* regulon (CysM, CysN, CysD and BCAL2684) were also increased in abundance in the HACs relative to the NACs (**Supplemental Table 4**). In contrast, proteins associated with cysteine catabolism such as IscS, a cysteine desulferase, were significantly decreased in abundance by ∼2 fold on day 7 and day 22 while GshA, a glutamate-cysteine ligase, was decreased in abundance by ∼3 fold on day 7 (**Supplemental Table 4**).

To examine whether the observed increased abundance of proteins involved in response to oxidative stress impacted the ability of *B. cenocepacia* to survive intracellularly, PBMC-derived macrophages from individuals with CF were infected with HACs or NACs at an MOI of 10:1 and the bacterial survival was determined after 24 h (**Figure 4A**). Proliferation of HACs within macrophages was significantly higher than that of NACs, with 25% more CFU in 24 hours relative to only 9% for NACs, thus showing a significant ∼2.8 fold increase in intra-macrophage survival in comparison to the NACs (p<0.05). This is comparable with the increased survival in CF-PBMCs observed by the late infection isolate, P1L compared to P1E (which had been isolated from the same patient 61 months earlier) (p<0.05), suggesting that long term exposure to hypoxia is contributing to enhanced survival of *B. cenocepacia* in CF macrophages.

**Figure 4.**
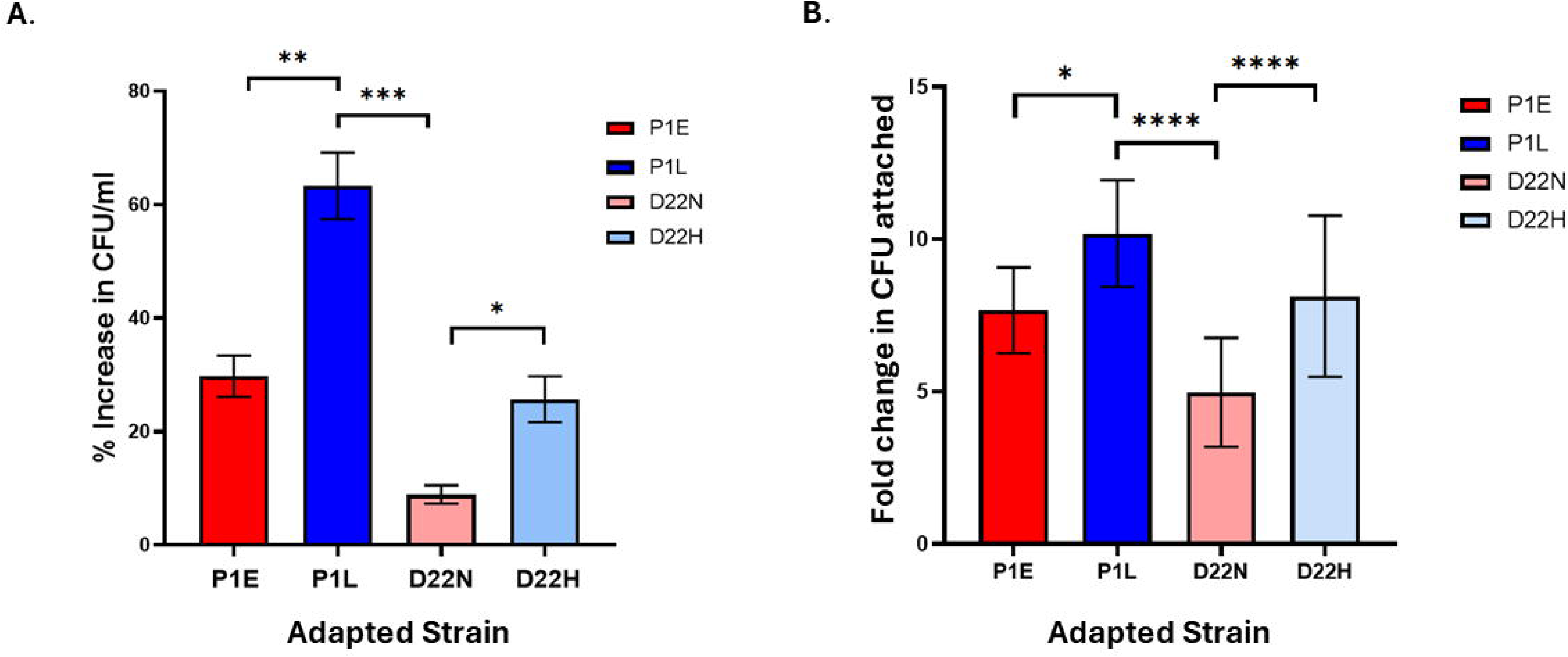
A) Effects of long-term exposure to hypoxia on survival in CF PBMC-derived macrophages. Mean survival of the early chronic infection isolate (P1E), the late chronic infection isolate (P1L) and 22-day populations of normoxia-adapted (D22N) and hypoxia-adapted *B. cenocepacia* cultures (D22H) in CF PBMC-derived macrophages 24 hours post-infection as determined by % increase in CFU/ml relative to CFU internalised at time zero, following inoculation with 5 x 10^5^ CFU/well in two independent experiments (and in duplicate). Error bars represent standard error of the mean. Statistically significant differences determined by one-way ANOVA analysis, shown as *p<0.05; **p<0.01; ***p<0.005 P1E: early infection isolate; P1L: late chronic infection isolate; D22N: 22-day normoxia-adapted cultures; D22H: 22-day hypoxia-adapted cultures. **B) Effects of long-term exposure to hypoxia on CF epithelial cell attachment.** Mean CFU attached to CFBE41o− cells determined by three independent experiments. Error bars represent standard error of the mean. Statistically significant differences determined by one-way ANOVA analysis shown as *p<0.05; ****p<0.001. P1E: early infection isolate; P1L: late chronic infection isolate; D22N: 22-day normoxia-adapted cultures; D22H: 22-day hypoxia-adapted cultures.

### Attachment to CF Lung Cells Increased in HACs

Four proteins implicated in host-cell attachment were significantly increased in abundance in the proteome of the HACs relative to the NACs (**Table 2**). The haemagglutinin-related autotransporter protein (BCAM0219) was consistently increased in abundance over the 22 day period in HACs by up to 840 fold relative to the NACs. Additionally, outer membrane protein A (BCAM0220) was also significantly increased in abundance in HACs by ∼660 fold on day 7 and by ∼13 fold on day 14. Interestingly, lectin protein (bclA) and fimbrial chaperone protein (BCAL1679) were unique to HACs on days 7 and 22 respectively. This aligns with our previous finding that late *B. cenocepacia* infection isolates showed increase attachment to host cells relative to early infection counterparts [22] and consequently we wanted to evaluate whether adaptation to hypoxia might drive this adaptation. CFBE41o-cells were inoculated with day 22 HACs or NACs and host-cell attachment determined by microbiological plating. Consistent with late infection isolates, the attachment of HACs to the CFBE41o-cells was ∼1.5 fold higher (p<0.005) than the NACs (**Figure 4B**), confirming that exposure to 6% oxygen had a significant effect on host cell attachment, comparable to that of persistent infection.

### Effect of Hypoxia exposure on *B. cenocepacia* biofilm formation

Although *B. cenocepacia* strains form strong biofilms *in vitro*, the importance of biofilm formation in chronic Bcc colonisation is not clear as Bcc biofilms have not been identified in explanted CF lungs to date [26]. Nevertheless, comparison of biofilm formation in P1E, P1L, NACs and HACs showed there were significant differences in biofilm formation at 48 hours (**Figure 5A**) as the early infection isolate P1E produced most biofilm, which was ∼1.8 fold more than the biofilms form by HACs (p<0.05).

**Figure 5.**
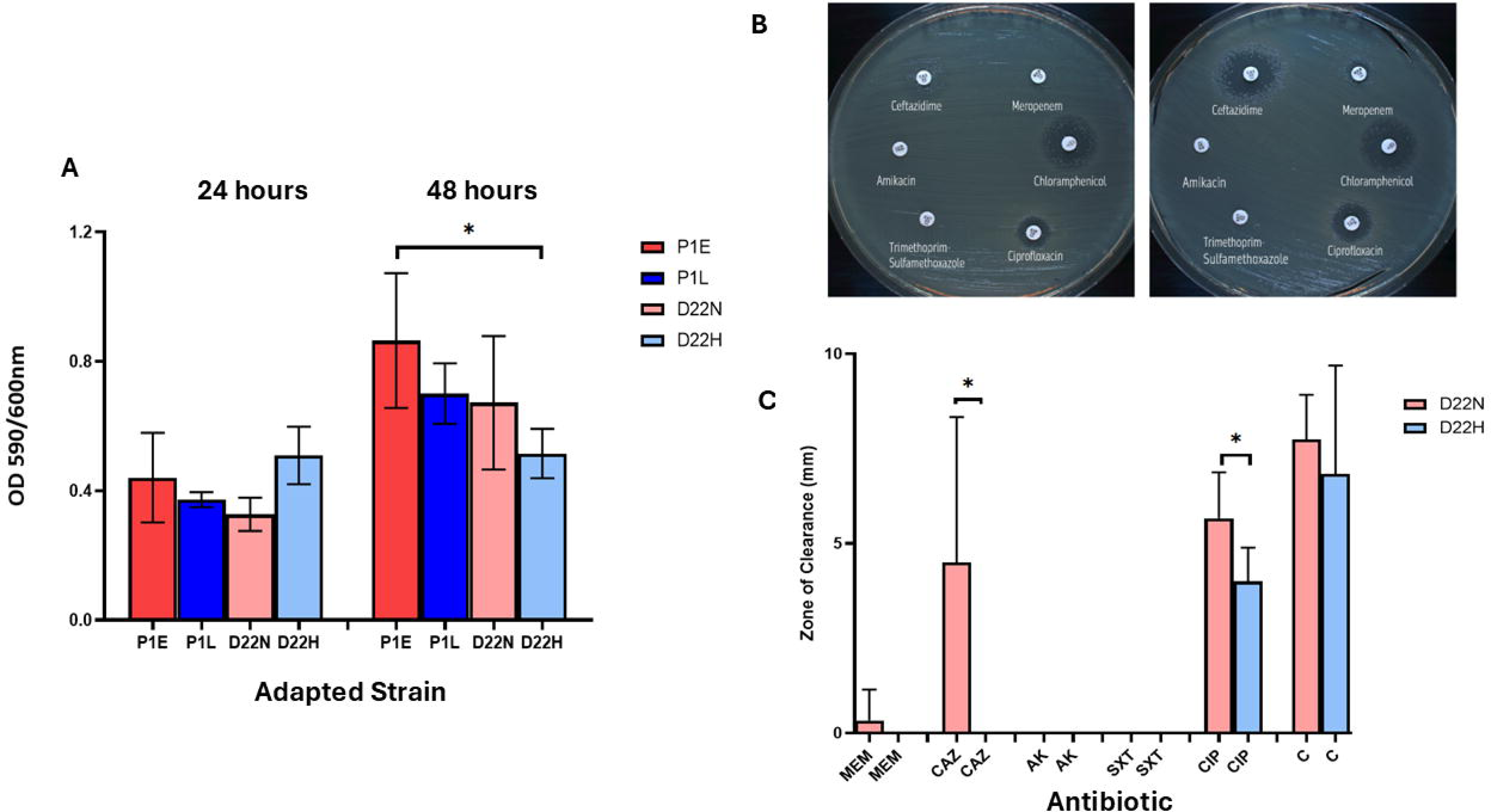
Effects of long-term exposure to hypoxia on biofilm formation and antibiotic susceptibility. **A)** Biofilm formation measured by final crystal violet staining (590nm/600nm) at 24-hours and 48-hours. Graph displays the mean biofilm formation of the early infection isolate (P1E), late chronic infection isolate (P1L) and pooled 22-day populations of normoxia-adapted (D22N) and hypoxia-adapted (D22H) *B. cenocepacia* cultures over three independent experiments. Error bars represent standard error of the mean. Statistically significant differences determined by one-way ANOVA analysis, shown as *p<0.05. P1E: early infection isolate; P1L: late chronic infection isolate; D22N: 22-day normoxia-adapted cultures; D22H: 22-day hypoxia-adapted cultures. **B)** Antibiotic susceptibility testing using disc diffusion assays on a panel of antibiotics against day-22 hypoxia-adapted *B. cenocepacia* cultures (left) compared to day-22 normoxia-adapted *B. cenocepacia* cultures (right) (n=3). **C)** Mean antibiotic susceptibility of 22-day populations of normoxia-adapted (D22N) and hypoxia-adapted cultures (D22H) to a panel of antibiotics. The data are representative of three independent experiments and the error bars represent the standard deviation. Statistical significance was determined by a paired t-test, shown as *p<0.05. Meropenem (MEM), Ceftazidime (CAZ), Amikacin (AK), Trimethoprim-Sulfamethoxazole (SXT), Ciprofloxacin (CIP), Chloramphenicol (C) (n=3).

### Long term Exposure to Hypoxia Enhances Antibiotic Resistance

Comparison of the proteomes of the HACs and P1E highlighted that regulatory proteins, transporter proteins and proteins implicated in cell wall formation were significantly changed in abundance in the proteome of the HACs relative to the early infection isolate (**Table 1**). A TetR family regulatory protein (BCAM2652) was unique to all three the HACs, while a MarR regulatory protein and an ABC transporter protein (BCAS0113) were significantly increased in abundance by ∼1.95 fold and ∼5.58 fold, respectively. MurF was also increased in abundance by ∼2.5 fold in the proteome of the HACs relative to early infection isolate, P1E (**Table 1).** Moreover, several proteins implicated in antibiotic resistance were also significantly increased in abundance over the course of 22-days in the HACs relative to the NACs (**Table 2**). Three of these proteins were unique to the HACs: an ABC transporter protein, a BceRS two-component system related protein and an acetyltransferase [27]. The acetyltransferase (BCAM1840) includes a Gcn5-related N-acetyltransferases (GNAT) domain often implicated in resistance to aminoglycosides [28]. Furthermore, a metallo-β-lactamase (BCAL0673) was significantly increased in abundance by ∼10 fold on day 14 (**Table 2**).

Consequently, to investigate whether adaptation to hypoxia contributes to antibiotic resistance, antibiotic susceptibility of the day 22 HACs and NACs to the clinically relevant antibiotics: amikacin, ceftazidime, chloramphenicol, ciprofloxacin, trimethoprim/ sulfamethoxazole and meropenem was determined (**Figure 5B)**. HACs demonstrated significantly lower susceptibility to ceftazidime (p<0.05) and ciprofloxacin (p<0.05) (**Figure 5C**) relative to the NACs. Both HACs and NACs were resistant to amikacin and trimethoprim/sulfamethoxazole.

### Long-term hypoxia culture promotes increased protease activity

Several proteases and proteins involved in protein turnover were identified as having changed abundance in HACs (**Supplemental Table 1 and 2**). Moreover, we had previously identified that late infection isolates showed increased abundance of multiple proteases [22]. Consequently, to examine whether adaptation to hypoxia drives increased protease activity, the HACs and NACs were inoculated into wells of milk-agar plates and protease activity was measured (**Figure 6A**). Consistent with our previous proteome analysis, the late infection isolates, P1L showed substantially more protease activity than the earlier isolate (**Figure 6B**). In addition, the HACs demonstrated significantly increased levels of protease activity compared with the NACs (p<0.001) and a ∼10-fold increase relative to the ancestral strain, P1E (p<0.001) (**Figure 6B**). To further examine the protease profiles of the clinical isolates and adapted cultures, the concentrated supernatants from each culture were analysed on zymograms (**Figure 6C**). The zymogram profile of the late infection isolate was quite distinct from the early infection, with additional bands of clearance at around 50kDa and ∼20kD in addition to the band at 33kDa. The zymography profiles of hypoxia-adapted cultures H1 and H3 also showed additional bands that were comparable to those of the late infection isolate P1L. Meanwhile, the protease profiles of the NACs were comparable to P1E, indicating hypoxia may be driving a similar change in protease expression to that observed in the chronic infection isolate. Interestingly, a subfamily M20A metallopeptidase (BCAM2149) is an extracellular peptidase that is implicated in the degradation of proteins in the host environment and was significantly increased in abundance by ∼6.5 fold on day 14 and ∼4 fold on day 22 in the proteome of the HACs relative to the NACs (**Table 2**). The molecular weight (MW) of BCAM2149 is 52.3kDa (**Figure 6C**), a band in this region is present in the zymograms for H1, H3 and P1L.

**Figure 6.**
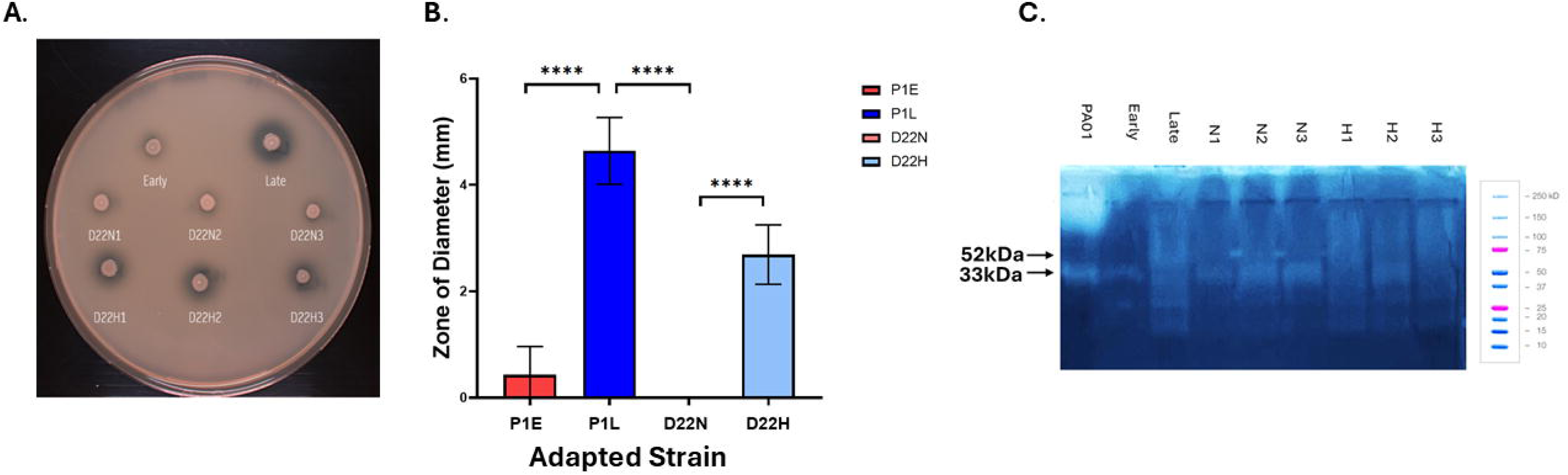
Effects of long-term exposure to hypoxia on protease activity. **A)** Protease activity of an early chronic infection isolate, late chronic infection isolate and 22-day populations of normoxia-adapted and hypoxia-adapted cultures presented as zones of clearance on a milk-agar plate. Image representative of three independent experiments. **B)** Mean protease activity of the early chronic infection isolate, late chronic infection isolate and grouped 22-day populations of normoxia-adapted (D22N) and hypoxia-adapted (D22H) *B. cenocepacia* cultures. The data represent the mean and standard deviation of three independent experiments. Statistical significance was determined by a one-way ANOVA analysis, shown as ****p<0.0001. **C)** Zymogram displaying the protease profile of an early chronic infection isolate, late chronic infection isolate and 22-day populations of normoxia-adapted and hypoxia-adapted cultures PA01 used as a positive control with LasB (33kDa) acting as a MW marker.

### Response to long-term hypoxia and Tryptophan Metabolism are linked

Numerous proteins associated with tryptophan catabolism were significantly changed in abundance in the HACs. Kynureninase (KynU) was significantly decreased in abundance by ∼4-to-5 fold on days 1, 7 and 22 in the HACs relative to the NACs (**Table 2**).Tryptophan 2,3-dioxygenase (KynA) was also significantly decreased in abundance by ∼33 to ∼57.5 fold on days 1 and 7 and by ∼13 fold at day 22. In addition, proteins associated with the tryptophan biosynthetic pathway were significantly increased in abundance in HACs, including tryptophan synthase alpha chain (TrpA), increased by ∼2.5 fold on day 14 and indole-3-glycerol phosphate synthase (TrpC), increased by ∼6 fold on day 14. These findings are significant because adaptation to hypoxia in Gram-negative pathogens has been associated with altered amino acid metabolism [29]. Tryptophan biosynthesis has recently been shown to play an important role in pathogenicity [30] and to further investigate the impact of these changes on tryptophan metabolism in HACs, the impact of tryptophan supplementation on growth was examined. Day 22 HACs showed significantly increased growth in minimal media supplemented with tryptophan relative to P1E (p<0.01) (**Figure 7A**) which mirrored the increased growth of the late infection isolate relative to the early infection isolate under the same conditions. This demonstrates that both P1L and HACs show altered tryptophan metabolism relative to the ancestral strain, P1E, and suggests that hypoxia exposure may drive changes in tryptophan metabolism. A number of proteins associated with the shikimate pathway, involved in aromatic amino acid biosynthesis [31], were also significantly increased in abundance in the proteome of the HACs relative to the ancestral strain **(Table 1),** including the phosphoenolpyruvate synthase regulatory protein (∼2 fold). Proteins in this pathway involved in the production of the chorismate were also increased in abundance in the HACs, including AroB (∼13 fold), AroC (∼3.5 fold), AroG (∼7 fold), chorismate mutase (∼10 fold), chorismate binding protein (∼3 fold) and isochorismatase (∼5.5 fold) **(Supplemental tables 3and 4)**.

**Figure 7.**
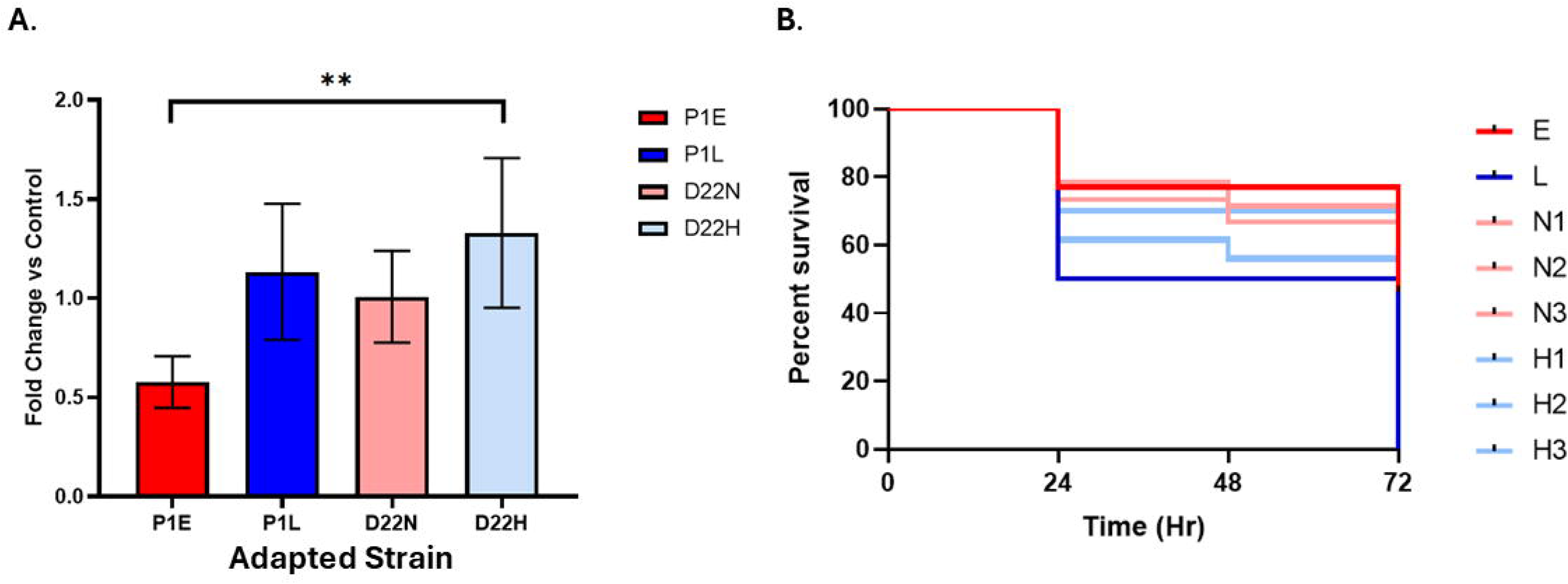
Effects of long-term exposure to hypoxia on growth in the presence of tryptophan and virulence in *G. mellonella*. **A)** Fold change in mean growth of P1E (E), P1L (L), D22 normoxia-adapted cultures (N) and D22 hypoxia-adapted cultures (H) *B. cenocepacia* cultures following overnight growth at 37°C in M9 media in the presence and absence of 100mM tryptophan supplementation. Data represent three independent experiments (6 replicates per condition per experiment). Error bars represent standard error of the mean. Statistically significant differences determined by one-way ANOVA analysis, shown as **p<0.01. **B)** Kaplan-Meier curve representative of the survival of *G. mellonella* infected with *B. cenocepacia* cultures over a period of 24-, 48-, 72-hours. E: early chronic infection isolate; L: late chronic infection isolate; D22N1, N2, N3: 22-day normoxia-adapted cultures; D22H1, H2, H3: 22-day hypoxia-adapted cultures.

### Long term Exposure to Hypoxia is linked to Phenylacetic Acid Degradation

In the longitudinal proteome analysis, the phenylacetic acid degradation pathway transcriptional regulator (PaaR) was significantly increased in abundance ∼2-fold in HACs relative to P1E (**Table 1**). Similarly in the pairwise analysis, phenylacetic acid degradation protein, PaaA, was significantly increased in abundance by ∼3 to 4 fold on days 14 and 22; PaaB was increased by ∼3 fold on day 22 and PaaF was increased by ∼8 fold on day 14 (**Table 2**). In contrast, Phenylacetate-coenzyme A ligase (PaaK) was significantly decreased in abundance on day 1 by ∼11.5 fold, on day 7 by ∼12 fold and on day 22 by ∼2.5 fold. Additionally, enoyl-CoA hydratase (PaaG) was decreased by days 7 and 22 while 3-hydroxybutyryl-CoA dehydrogenase (PaaH) was decreased by 2 to 3 fold on days 1 and 7. These changes are of interest because phenylacetic acid degradation and reduced paaK expression is associated with enhanced virulence in *B. cenocepacia* [32].

Given the changes observed in proteins associated with virulence in HACs, *G. mellonella* larvae were used to examine the impact of these changes on virulence *in vivo*. P1L was much more virulent than P1E, with 50% larval death at 24 h compared to 80% in P1E inoculated larvae, (**Figure 7B**). Interestingly, the HACs mediated the killing of more larvae than NACs overall, with flask 1 of the HACs being the most virulent out of the adapted cultures across three independent experiments.

## Discussion

The genetic plasticity of *B. cenocepacia* facilitates efficient adaptation to a range of hostile environments [13]. The ability to thrive in adverse conditions including disinfectants, biocides, pollutants, crude oil and personal care products is a hallmark of *Bcc* [33]. Adaptation to stress is critical for colonisation in the CF lung, and in particular, sensing and responding to low oxygen is key to survival and proliferation [34]. We have shown that exposure to hypoxia alone over a period of 22 days, increased host cell attachment, intramacrophage survival, protease activity and resistance to both ciprofloxacin and ceftazidime.

We had previously reported that proteomes of two late infection isolates had changed over time of infection and consequently, we focussed on the impact of hypoxia exposure on the proteome directly. The proteome of HACs was also dramatically altered following exposure to hypoxia for 22 days. Thus, in order to probe the relationship between adaptation following exposure to long-term hypoxia and changes identified in chronic infection isolates, we compared the alterations in the proteome of the HACs over the 22-day period with those that we had identified as being altered in the late infection isolate P1L (relative to its earlier counterpart (P1E)) [20] **(Table 3)**. It is noteworthy that the 39 proteins showing significantly increased in abundance were common to the proteomes of both the HACs (relative to NACs) and the late infection isolate, P1L, (relative to P1E), suggesting a shared response mechanism. It was also found that the same 42 proteins were significantly decreased in abundance in both proteomes. Interestingly, 8 of these common proteins that showed increased in abundance were encoded on the Lxa locus (**Table 3**). Of these proteins, there were three USPs, a HSP, a phospholipid binding protein (PBP), a phasin family protein, acetoacetyl-CoA reductase and a DUF4426-domain containing protein. FixK was consistently increased in abundance in HACs and in the proteome of the late chronic infection isolate. FixK is regulated by the oxygen-sensing two component system FixLJ, which is positively selected for during chronic infection [35]. A recent review of several studies suggests that FixK, may be involved in the regulation of the Lxa locus [7], but this has not been proven directly.

**Table 3.**
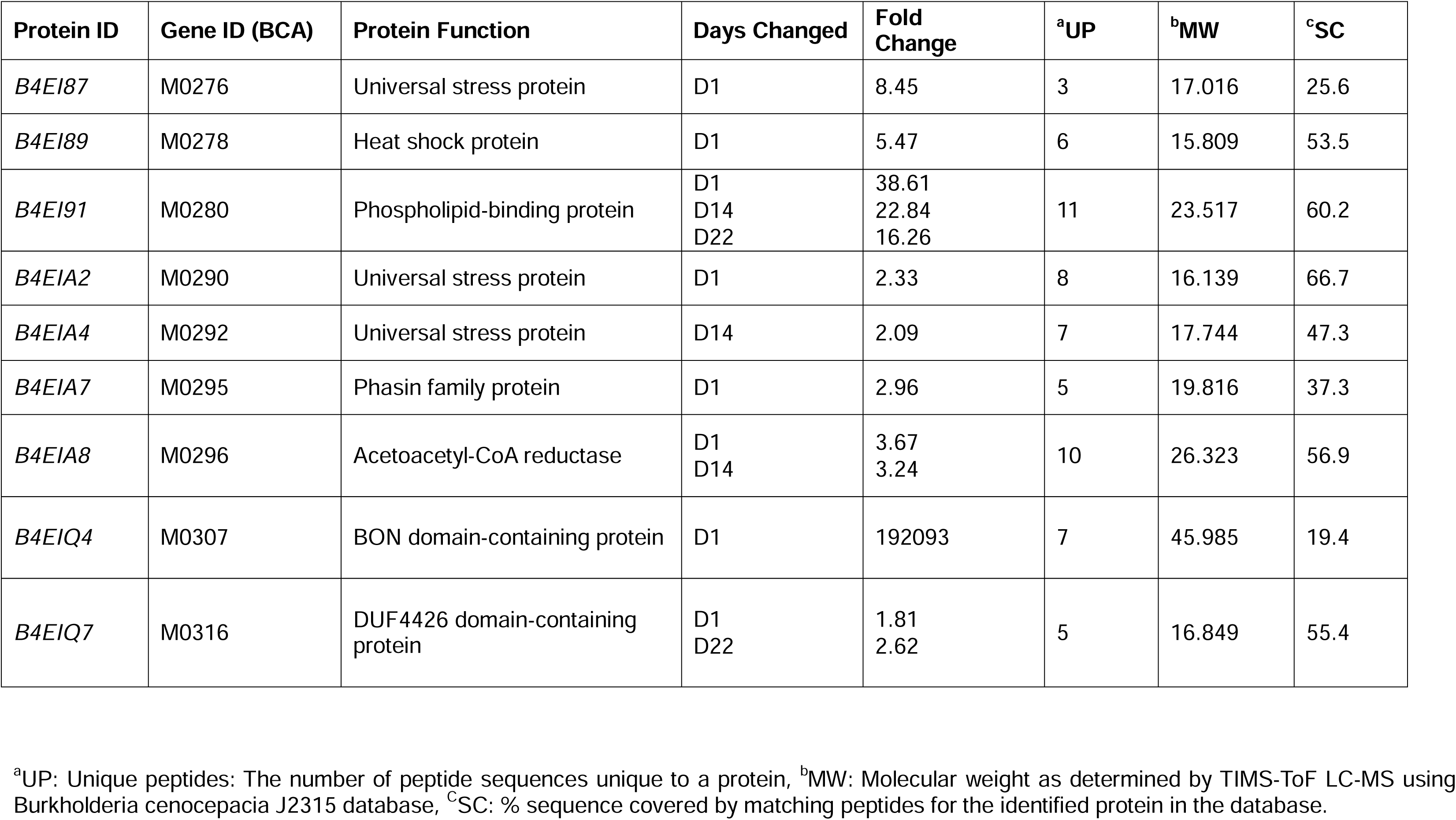
Lxa-encoded proteins significantly changed in abundance following adaptation to hypoxia and previously altered in abundance during chronic infection.

| Protein ID | Gene ID (BCA) | Protein Function | Days Changed | Fold Change | <sup>a</sup>UP | <sup>b</sup>MW | <sup>c</sup>SC |
| --- | --- | --- | --- | --- | --- | --- | --- |
| *B4EI87* | M0276 | Universal stress protein | D1 | 8.45 | 3 | 17.016 | 25.6 |
| *B4EI89* | M0278 | Heat shock protein | D1 | 5.47 | 6 | 15.809 | 53.5 |
| *B4EI91* | M0280 | Phospholipid-binding protein | D1<br>D14<br>D22 | 38.61<br>22.84<br>16.26 | 11 | 23.517 | 60.2 |
| *B4EIA2* | M0290 | Universal stress protein | D1 | 2.33 | 8 | 16.139 | 66.7 |
| *B4EIA4* | M0292 | Universal stress protein | D14 | 2.09 | 7 | 17.744 | 47.3 |
| *B4EIA7* | M0295 | Phasin family protein | D1 | 2.96 | 5 | 19.816 | 37.3 |
| *B4EIA8* | M0296 | Acetoacetyl-CoA reductase | D1<br>D14 | 3.67<br>3.24 | 10 | 26.323 | 56.9 |
| *B4EIQ4* | M0307 | BON domain-containing protein | D1 | 192093 | 7 | 45.985 | 19.4 |
| *B4EIQ7* | M0316 | DUF4426 domain-containing protein | D1<br>D22 | 1.81<br>2.62 | 5 | 16.849 | 55.4 |

<sup>a</sup>UP: Unique peptides: The number of peptide sequences unique to a protein, <sup>b</sup>MW: Molecular weight as determined by TIMS-ToF LC-MS using *Burkholderia cenocepacia* J2315 database, <sup>c</sup>SC: % sequence covered by matching peptides for the identified protein in the database.

Although no significant increase was observed in the abundance Lxa locus-encoded metallo-β-lactamase BCAM0300, in HACs (previously shown to be increased in abundance in late infection isolates) [20], another metallo-β-lactamase (BCAL0673) was significantly increased in abundance in the HACs relative to the NACs. This coupled with the finding that all three HACs displayed significantly increased resistance to ceftazidime indicates that hypoxia exposure alone may drive the expression of antibiotic resistance. The response to environmental stress can also induce morphological changes which may also confer resistance to cell wall-targeting β-lactams [36]. SCVs are clinically relevant morphotypes that are implicated in a persistent lifestyle [37]. SCVs are also associated with increased antibiotic resistance, enhanced biofilm formation and elevated levels of stress tolerance [36]. Overall, this highlights the role of hypoxia in driving increased antibiotic resistance in *B. cenocepacia*.

Long-term hypoxia drives *gyrB* mutations in *S. aureus* [38], and consistent with this, HACs showed reduced susceptibility to ciprofloxacin, a fluoroquinolone that targets DNA gyrase. The efficacy of ciprofloxacin is influenced by the conformational state of DNA gyrase [39]. It is possible that long-term adaptation to hypoxia has stably remodelled nucleoid architecture [40]. In our proteome analysis, we saw changes in the abundance of proteins associated with regulating the conformational state of the nucleoid, for example ParC and ParE, which are significantly decreased in abundance in the HACs relative to P1E. Interestingly, a H-NS protein was found to be unique to normoxia (i.e. absent in HACs), further suggesting that long-term exposure to low oxygen is driving DNA remodelling. Overall, these data highlight the impact of hypoxia on the differential transcription of genes in order to facilitate adaptation to low oxygen.

Survival in stressful conditions is often associated with a transition to a more persistent state [41]. In *P. aeruginosa,* chronic CF infection is commonly associated with mucoidy and increased biofilm formation [42]. In contrast, the *B. cenocepacia* late chronic infection isolates showed reduced levels of biofilm formation relative to the early infection isolate. Tryptophan availability has been shown to inhibit biofilm formation and enhance virulence [43, 44]. A consistent decrease in proteins associated with tryptophan degradation and a consistent increase in proteins associated with tryptophan biosynthesis were observed the HACs. Che et al. (2025) revealed that tryptophan synthase alpha chain (*trpA)* mutants showed significantly attenuated virulence, with a reduction in tryptophan levels, biofilm formation, motility, hemolysis and quorum sensing highlighting the importance of tryptophan in *Vibrio anguillarum* virulence [30]. An increase in intracellular tryptophan and other aromatic amino acids following long-term exposure to hypoxia may arise from activity of the shikimate pathway (**Figure 8**) [31]. Decreased phenylacetic acid (PAA) levels are associated with enhanced QS communication [45]. Consistent increases in the abundance of proteins associated with phenylalanine degradation and decreases in the abundance of PaaK were also observed in the HACs, which are associated with increased activity of the CepIR system [32]. The CepIR is implicated in the increased expression of virulence factors and reduced intracellular c-di-GMP levels, and its expression was previously shown to be increased under short term low oxygen [19]. Overall, this suggests that adaptation to hypoxia increases the availability of amino acids important in chronic infection.

**Figure 8.**
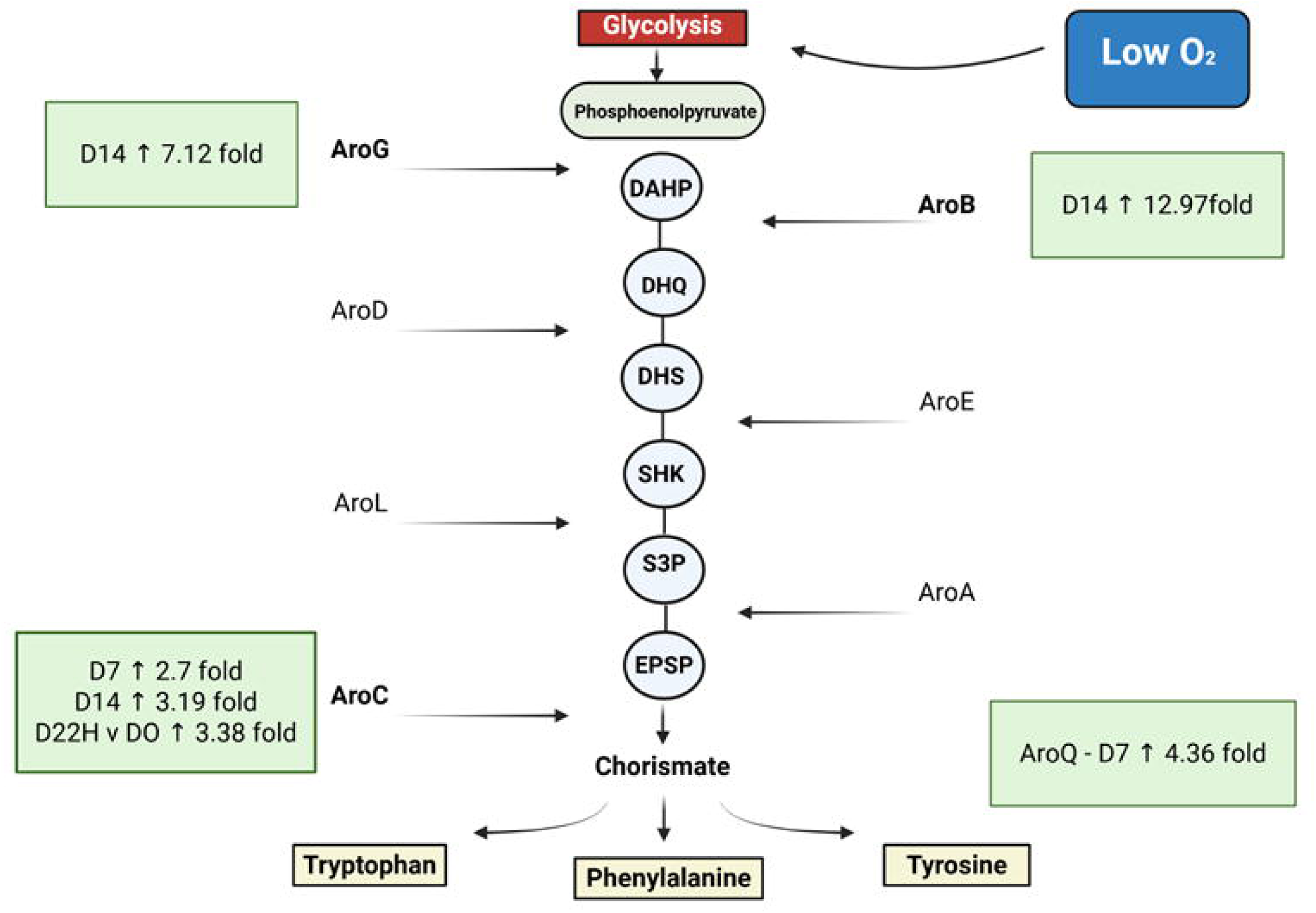
Schematic illustrating proteins associated with Shikimate Pathway. Diagram highlights the proteins significantly changed in abundance in the proteome of the hypoxia-adapted cultures relative to the normoxia-adapted cultures and the early infection isolate: AroG, AroB, AroC and AroQ. DAHP: 3-Deoxy-D-arabinoheptulosonate 7-phosphate, DHQ: 3-dehydroquinate, DHS: 3-dehydroshikimate, SHK: Shikimate, SP3: Shikimate-3-Phosphate, EPSP: 5-Enolpyruvylshikimate-3-phosphate.

*B. cenocepacia* expresses lectins, fimbriae and type IV cable pili to attach to the host epithelium [46, 47]. HACs showed more attachment to CFBE cells than the NACs which may have been due, a least in part, to the presence of the lectin (BclA) and fimbrial chaperone protein (BCAL1679) in HACs (and not in NACs). BclA was previously shown to be significantly upregulated in response to short-term exposure to low oxygen conditions [19]. The concomitant increases in host-cell attachment evident HACs and PIL relative to NACs and P1E respectively, again suggests that hypoxia plays a role in enhancing host-cell attachment and ultimately, in the increased host-cell attachment previously observed over time of chronic infection [20].

*Bcc* also secretes metalloproteases to degrade proteins and tissues to support colonisation [48]. An increased abundance of serine proteases, cysteine peptidases and metalloproteases were observed in HACs. In particular, BCAM2149 is an M20 subfamily carboxypeptidase, a homolog of which has been shown to be essential for host tissue invasion in *Fransicella tularnesis* [49]. As this metallopeptidase and phospholipase C were consistently increased in abundance in HACs, it is possible that hypoxia is driving tissue invasion.

To establish chronic infection, *B. cenocepacia* must also be able to tolerate oxidative stress. A hallmark of chronic *B. cenocepacia* infection is the capacity to survive and proliferate within macrophages [50]. The CF lung has intrinsically high levels of ROS [51]. Proteins associated with oxidative stress e.g. catalase, were significantly increased in abundance in the proteome of P1L relative to P1E, indicating that tolerance to ROS may be critical for chronic infection [20]. Interestingly, many of the same proteins are increased in abundance in the proteome of the HACs, such as SodB, HmpA, AhpC, AhpD, suggesting long-term hypoxia may be driving increased tolerance to oxidative stress and ultimately enhancing intracellular survival.

## Conclusions

In conclusion, adaptation to hypoxia drives proteome and phenotypic changes that are consistent with the changes observed in chronic infection isolates. This suggests that adaptation to low oxygen may prime a pathogen for chronic infection. Intensive treatment with oxygen may help deter the development of chronic infection by reducing the opportunity to adapt to low oxygen levels and recent studies have shown benefits of hyperbaric oxygen in clearance of *P. aeruginosa*. In the longer term, better understanding of the underlying mechanisms, may lead to the development of novel therapeutics that target these oxygen sensing and response systems. For example, development of a therapy that inhibits the FixK activity, such as those that repress the FixLJ two component system [52] could potentially dampen the hypoxic response and hinder the downstream expression of traits that lead to the development of chronic infection.

## Data Availability

The mass spectrometry proteomics data have been deposited to the ProteomeXchange Consortium via the PRIDE partner repository with the reference: 1-20250806-135227-3733916

## List of Abbreviations

CF: Cystic Fibrosis
CFTR: Cystic Fibrosis Transmembrane Conductance Regulator
Bcc: Burkholderia cepacia complex
LB: Lysogeny Broth
MEM: Minimal essential media
BSA: Bovine Serum Albumin
SCV: Small Colony Variant
HAC: Hypoxia-adapted Culture
NAC: Normoxia-adapted Culture
ROS: Reactive Oxygen Species
USP: Universal Stress Protein
HSP: Heat Shock Protein
CSP: Cold Shock Protein
PBMC: Peripheral Blood Mononuclear Cell
CFBE41o-: Cystic Fibrosis Bronchial Epithelial (cell line)
GNAT: Gcn5-related N-acetyltransferase
PAA: Phenylacetic Acid
P1E: Early infection isolate of *B. cenocepacia*
P1L: Late infection isolate of *B. cenocepacia*
GO: Gene Ontology

## Bibliography

1. Fanen P, Wohlhuter-Haddad A, Hinzpeter A: Genetics of cystic fibrosis: CFTR mutation classifications toward genotype-based CF therapies. Int J Biochem Cell Biol 2014, 52:94–102.

2. Leclair LW, Hogan DA: Mixed bacterial-fungal infections in the CF respiratory tract. Med Mycol 2010, 48 Suppl 1:S125–132.

3. Blanchard AC, Waters VJ: Opportunistic Pathogens in Cystic Fibrosis: Epidemiology and Pathogenesis of Lung Infection. J Pediatric Infect Dis Soc 2022, 11(Supplement_2):S3–s12.

4. Dobrindt U, Zdziarski J, Salvador E, Hacker J: Bacterial genome plasticity and its impact on adaptation during persistent infection. Int J Med Microbiol 2010, 300(6):363–366.

5. Dawan J, Ahn J: Bacterial Stress Responses as Potential Targets in Overcoming Antibiotic Resistance. Microorganisms 2022, 10(7).

6. Poole K: Bacterial stress responses as determinants of antimicrobial resistance. J Antimicrob Chemother 2012, 67(9):2069–2089.

7. Carey CJ, Duggan N, Drabinska J, McClean S: Harnessing hypoxia: bacterial adaptation and chronic infection in cystic fibrosis. FEMS Microbiol Rev 2025, 49.

8. Jurado-Martin I, Sainz-Mejias M, McClean S: Pseudomonas aeruginosa: An Audacious Pathogen with an Adaptable Arsenal of Virulence Factors. Int J Mol Sci 2021, 22(6).

9. Velez LS, Aburjaile FF, Farias ARG, Baia ADB, Oliveira WJ, Silva AMF, Benko-Iseppon AM, Azevedo V, Brenig B, Ham JH et al: Burkholderia semiarida sp. nov. and Burkholderia sola sp. nov., two novel B. cepacia complex species causing onion sour skin. Syst Appl Microbiol 2023, 46(3):126415.

10. Drevinek P, Mahenthiralingam E: Burkholderia cenocepacia in cystic fibrosis: epidemiology and molecular mechanisms of virulence. Clin Microbiol Infect 2010, 16(7):821–830.

11. Scoffone VC, Chiarelli LR, Trespidi G, Mentasti M, Riccardi G, Buroni S: Burkholderia cenocepacia Infections in Cystic Fibrosis Patients: Drug Resistance and Therapeutic Approaches. Front Microbiol 2017, 8:1592.

12. O’Grady EP, Sokol PA: Burkholderia cenocepacia differential gene expression during host-pathogen interactions and adaptation to the host environment. Front Cell Infect Microbiol 2011, 1:15.

13. Mahenthiralingam E, Urban TA, Goldberg JB: The multifarious, multireplicon Burkholderia cepacia complex. Nat Rev Microbiol 2005, 3(2):144–156.

14. Courtney JM, Dunbar KE, McDowell A, Moore JE, Warke TJ, Stevenson M, Elborn JS: Clinical outcome of Burkholderia cepacia complex infection in cystic fibrosis adults. J Cyst Fibros 2004, 3(2):93–98.

15. Blackburn L, Brownlee K, Conway S, Denton M: ’Cepacia syndrome’ with Burkholderia multivorans, 9 years after initial colonization. J Cyst Fibros 2004, 3(2):133–134.

16. Greenwald MA, Wolfgang MC: The changing landscape of the cystic fibrosis lung environment: From the perspective of Pseudomonas aeruginosa. Curr Opin Pharmacol 2022, 65:102262.

17. André AC, Laborde M, Marteyn BS: The battle for oxygen during bacterial and fungal infections. Trends Microbiol 2022, 30(7):643–653.

18. Filkins LM, O’Toole GA: Cystic Fibrosis Lung Infections: Polymicrobial, Complex, and Hard to Treat. PLoS Pathog 2015, 11(12):e1005258.

19. Sass AM, Schmerk C, Agnoli K, Norville PJ, Eberl L, Valvano MA, Mahenthiralingam E: The unexpected discovery of a novel low-oxygen-activated locus for the anoxic persistence of Burkholderia cenocepacia. Isme J 2013, 7(8):1568–1581.

20. Cullen L, O’Connor A, McCormack S, Owens RA, Holt GS, Collins C, Callaghan M, Doyle S, Smith D, Schaffer K et al: The involvement of the low-oxygen-activated locus of Burkholderia cenocepacia in adaptation during cystic fibrosis infection. Sci Rep 2018, 8(1):13386.

21. O’Connor A, Jurado-Martin I, Mysior MM, Manzira AL, Drabinska J, Simpson JC, Lucey M, Schaffer K, Berisio R, McClean S: A universal stress protein upregulated by hypoxia has a role in Burkholderia cenocepacia intramacrophage survival: Implications for chronic infection in cystic fibrosis. Microbiologyopen 2023, 12(1):e1311.

22. Cullen L, O’Connor A, Drevinek P, Schaffer K, McClean S: Sequential Burkholderia cenocepacia Isolates from Siblings with Cystic Fibrosis Show Increased Lung Cell Attachment. American journal of respiratory and critical care medicine 2017, 195(6):832–835.

23. Tikhomirova A, Rahman MM, Kidd SP, Ferrero RL, Roujeinikova A: Cysteine and resistance to oxidative stress: implications for virulence and antibiotic resistance. Trends Microbiol 2024, 32(1):93–104.

24. Chung JW, Speert DP: Proteomic identification and characterization of bacterial factors associated with Burkholderia cenocepacia survival in a murine host. Microbiology (Reading) 2007, 153(Pt 1):206–214.

25. Cobley JN: 50 shades of oxidative stress: A state-specific cysteine redox pattern hypothesis. Redox Biol 2023, 67:102936.

26. Schwab U, Abdullah LH, Perlmutt OS, Albert D, Davis CW, Arnold RR, Yankaskas JR, Gilligan P, Neubauer H, Randell SH et al: Localization of Burkholderia cepacia complex bacteria in cystic fibrosis lungs and interactions with Pseudomonas aeruginosa in hypoxic mucus. Infect Immun 2014, 82(11):4729–4745.

27. Kawada-Matsuo M, Yoshida Y, Nakamura N, Komatsuzawa H: Role of two-component systems in the resistance of Staphylococcus aureus to antibacterial agents. Virulence 2011, 2(5):427–430.

28. Czub MP, Zhang B, Chiarelli MP, Majorek KA, Joe L, Porebski PJ, Revilla A, Wu W, Becker DP, Minor W et al: A Gcn5-Related N-Acetyltransferase (GNAT) Capable of Acetylating Polymyxin B and Colistin Antibiotics in Vitro. Biochemistry 2018, 57(51):7011–7020.

29. Bange G, Brodersen DE, Liuzzi A, Steinchen W: Two P or Not Two P: Understanding Regulation by the Bacterial Second Messengers (p)ppGpp. Annu Rev Microbiol 2021, 75:383–406.

30. Che J, Liu B, Fang Q, Nissa MU, Luo T, Wang L, Bao B: Biological studies reveal the role of trpA gene in biofilm formation, motility, hemolysis and virulence in Vibrio anguillarum. Microb Pathog 2025, 200:107331.

31. Herrmann KM, Weaver LM: THE SHIKIMATE PATHWAY. Annu Rev Plant Physiol Plant Mol Biol 1999, 50:473–503.

32. Lightly TJ, Frejuk KL, Groleau MC, Chiarelli LR, Ras C, Buroni S, Déziel E, Sorensen JL, Cardona ST: Phenylacetyl Coenzyme A, Not Phenylacetic Acid, Attenuates CepIR-Regulated Virulence in Burkholderia cenocepacia. Appl Environ Microbiol 2019, 85(24).

33. Tavares M, Kozak M, Balola A, Sá-Correia I: Burkholderia cepacia Complex Bacteria: a Feared Contamination Risk in Water-Based Pharmaceutical Products. Clin Microbiol Rev 2020, 33(3).

34. Taabazuing CY, Hangasky JA, Knapp MJ: Oxygen sensing strategies in mammals and bacteria. J Inorg Biochem 2014, 133:63–72.

35. Schaefers MM, Liao TL, Boisvert NM, Roux D, Yoder-Himes D, Priebe GP: An Oxygen-Sensing Two-Component System in the Burkholderia cepacia Complex Regulates Biofilm, Intracellular Invasion, and Pathogenicity. PLoS pathogens 2017, 13(1):e1006116.

36. Shen JP, Chou CF: Morphological plasticity of bacteria-Open questions. Biomicrofluidics 2016, 10(3):031501.

37. Starkey M, Hickman JH, Ma L, Zhang N, De Long S, Hinz A, Palacios S, Manoil C, Kirisits MJ, Starner TD et al: Pseudomonas aeruginosa rugose small-colony variants have adaptations that likely promote persistence in the cystic fibrosis lung. J Bacteriol 2009, 191(11):3492–3503.

38. Hull RC, Wright RCT, Sayers JR, Sutton JAF, Rzaska J, Foster SJ, Brockhurst MA, Condliffe AM: Antibiotics Limit Adaptation of Drug-Resistant Staphylococcus aureus to Hypoxia. Antimicrob Agents Chemother 2022, 66(12):e0092622.

39. Sissi C, Perdonà E, Domenici E, Feriani A, Howells AJ, Maxwell A, Palumbo M: Ciprofloxacin affects conformational equilibria of DNA gyrase A in the presence of magnesium ions. J Mol Biol 2001, 311(1):195–203.

40. Hołówka J, Zakrzewska-Czerwińska J: Nucleoid Associated Proteins: The Small Organizers That Help to Cope With Stress. Front Microbiol 2020, 11:590.

41. Trastoy R, Manso T, Fernández-García L, Blasco L, Ambroa A, Pérez Del Molino ML, Bou G, García-Contreras R, Wood TK, Tomás M: Mechanisms of Bacterial Tolerance and Persistence in the Gastrointestinal and Respiratory Environments. Clin Microbiol Rev 2018, 31(4).

42. Pritt B, O’Brien L, Winn W: Mucoid Pseudomonas in cystic fibrosis. Am J Clin Pathol 2007, 128(1):32–34.

43. Brandenburg KS, Rodriguez KJ, McAnulty JF, Murphy CJ, Abbott NL, Schurr MJ, Czuprynski CJ: Tryptophan inhibits biofilm formation by Pseudomonas aeruginosa. Antimicrob Agents Chemother 2013, 57(4):1921–1925.

44. Schwager S, Agnoli K, Kothe M, Feldmann F, Givskov M, Carlier A, Eberl L: Identification of Burkholderia cenocepacia strain H111 virulence factors using nonmammalian infection hosts. Infect Immun 2013, 81(1):143–153.

45. Pribytkova T, Lightly TJ, Kumar B, Bernier SP, Sorensen JL, Surette MG, Cardona ST: The attenuated virulence of a Burkholderia cenocepacia paaABCDE mutant is due to inhibition of quorum sensing by release of phenylacetic acid. Mol Microbiol 2014, 94(3):522–536.

46. Tomich M, Mohr CD: Adherence and autoaggregation phenotypes of a Burkholderia cenocepacia cable pilus mutant. FEMS Microbiol Lett 2003, 228(2):287–297.

47. Marchetti R, Malinovska L, Lameignère E, Adamova L, de Castro C, Cioci G, Stanetty C, Kosma P, Molinaro A, Wimmerova M et al: Burkholderia cenocepacia lectin A binding to heptoses from the bacterial lipopolysaccharide. Glycobiology 2012, 22(10):1387–1398.

48. Kooi C, Corbett CR, Sokol PA: Functional analysis of the Burkholderia cenocepacia ZmpA metalloprotease. J Bacteriol 2005, 187(13):4421–4429.

49. Zellner B, Mengin-Lecreulx D, Tully B, Gunning WT, 3rd, Booth R, Huntley JF: A Francisella tularensis L,D-carboxypeptidase plays important roles in cell morphology, envelope integrity, and virulence. Mol Microbiol 2021, 115(6):1357–1378.

50. Mesureur J, Feliciano JR, Wagner N, Gomes MC, Zhang L, Blanco-Gonzalez M, van der Vaart M, O’Callaghan D, Meijer AH, Vergunst AC: Macrophages, but not neutrophils, are critical for proliferation of Burkholderia cenocepacia and ensuing host-damaging inflammation. PLoS Pathog 2017, 13(6):e1006437.

51. Moliteo E, Sciacca M, Palmeri A, Papale M, Manti S, Parisi GF, Leonardi S: Cystic Fibrosis and Oxidative Stress: The Role of CFTR. Molecules 2022, 27(16).

52. Mansour KE, Qi Y, Yan M, Ramstrom O, Priebe GP, Schaefers MM: Small-molecule activators of a bacterial signaling pathway inhibit virulence. bioRxiv 2023.

