## Supplemental Tables 1 and 2 for "Adaptation of *Burkholderia cenocepacia* to low oxygen drives changes consistent with adaptation to chronic infection"

**Supplementary Material**

**Supplementary Table 1.** Functional enrichment of proteins significantly changed in abundance in day 22 hypoxia-adapted cultures relative to the early infection isolate.

| **COG Categories** | **Number of Proteins** |
| --- | --- |
| Cell wall/membrane/envelope biogenesis | 33 |
| Defence mechanisms | 6 |
| Secondary metabolites biosynthesis, transport and catabolism | 4 |
| Amino acid transport and metabolism | 26 |
| Carbohydrate transport and metabolism | 12 |
| Cell cycle control, cell division, chromosome partitioning | 13 |
| Coenzyme transport and metabolism | 17 |
| Energy production and conversion | 31 |
| Function Unknown | 74 |
| Inorganic ion transport and metabolism | 9 |
| Intracellular trafficking, secretion, and vesicular transport | 7 |
| Lipid transport and metabolism | 12 |
| No COG Category Assigned | 17 |
| Nucleotide transport and metabolism | 23 |
| Post-translational modification, protein turnover, and chaperones | 21 |
| Replication, recombination and repair | 16 |
| Signal transduction mechanisms | 6 |
| Transcription | 19 |
| Translation, ribosomal structure and biogenesis | 18 |
| **Total** | **364** |

**Supplementary Table 2.** Functional enrichment of proteins significantly changed in abundance in day 22 hypoxia-adapted cultures relative to day 22 normoxia-adapted cultures.

| **COG Categories** | **Number of Proteins** |
| --- | --- |
| Translation, ribosomal structure and biogenesis | 28 |
| Transcription | 27 |
| Replication, recombination and repair | 13 |
| Cell cycle control, cell division, chromosome partitioning | 8 |
| Cell wall/membrane/envelope biogenesis | 21 |
| Cell motility | 1 |
| Post-translational modification, protein turnover, and chaperones | 14 |
| Signal transduction mechanisms | 9 |
| Intracellular trafficking, secretion, and vesicular transport | 4 |
| Defence mechanisms | 2 |
| Energy production and conversion | 45 |
| Amino acid transport and metabolism | 43 |
| Nucleotide transport and metabolism | 18 |
| Carbohydrate transport and metabolism | 23 |
| Coenzyme transport and metabolism | 19 |
| Lipid transport and metabolism | 25 |
| Inorganic ion transport and metabolism | 21 |
| Secondary metabolites biosynthesis, transport and catabolism | 15 |
| Function unknown | 59 |
| No COG category assigned | 8 |
| ***Total*** | **403** |
